# Contig-Resolved Genetic Context of Antimicrobial Resistome in Sediments of Two Contrasting Wastewater-Impacted Rivers

**DOI:** 10.64898/2026.11.03.763893

**Authors:** Arun Kashyap, Vikas Sonkar, Indumathi M. Nambi, Shashidhar Thatikonda

## Abstract

River sediments harbor diverse antimicrobial resistance (AMR) determinants; however, gene abundance alone provides limited insight into their genomic context, mobility potential, and associated bacterial hosts. This study employed an assembly- and read-mapping metagenomic framework to surface-sediment metagenomes (40.78 Gb clean data) from two contrasting wastewater-impacted Indian rivers, Adyar (Chennai) and Musi (Hyderabad), retaining sample-, contig- and open reading frame level provenance. ARG-carrying contigs (ACCs; n = 2,402) were annotated for mobile genetic elements (MGEs), biocide/metal resistance genes (BMRGs), virulence factor genes (VFGs), nitrogen-cycling genes (NCycGs), and putative host taxonomy. The resistomes exhibited substantial diversity (264 ARG subtypes) and spatial heterogeneity, with subtype richness in the Adyar (up to 171 subtypes) than in the Musi, where richness progressively declined downstream. Rank IV determinants dominated ARG abundance (77.82%), whereas Rank I determinants accounted for only 9.89%. MGEs were detected on just 3.12% (75/2,402) of ACCs, predominantly comprising plasmid-like contigs and proximal ARG-MGE associations. Co-occurrence with BMRGs was observed on 386 ACCs, whereas VFGs were uncommon (26 ACCs) and primarily represented dual-annotated efflux and envelope-integrity determinants rather than independent virulence loci. NCyc annotations were sparse (12 ACCs), with no evidence of complete nitrification or denitrification pathways. Taxonomic classification resolved 217 ACCs to the species level, with no matches to curated human pathogenic bacteria. Only two *Acinetobacter*-assigned ACCs exhibited both genus-level pathogenicity and mobility evidence. These findings highlight the importance of integrating mobility and host-taxonomic information to distinguish baseline environmental resistance from potential priorities for AMR surveillance.

## 1. Introduction

Antimicrobial resistance (AMR) is a major One Health challenge linking human, animal, and environmental microbial communities (Liguori et al. 2022; Larsson and Flach 2022). Antimicrobial use in human medicine, animal production, and agriculture exerts selective pressure that favors resistant microorganisms (Taneja and Sharma 2019; He et al. 2020; Musoke et al. 2021). In parallel, the release of antimicrobial residues, resistant microorganisms, and antibiotic resistance genes (ARGs) into the environment may facilitate their persistence and dissemination across interconnected environmental compartments (Kashyap et al. 2023; Sonkar et al. 2025). However, ARGs are not exclusively anthropogenic and occur naturally within indigenous microbial communities (Surette and Wright 2017; Klümper et al. 2024). The environmental resistome therefore comprises naturally occurring resistance functions as well as those enriched, redistributed, or mobilized under anthropogenic pressures. Aquatic systems are particularly relevant in this context because they integrate biological and chemical inputs from multiple sources across hydrologically connected catchments (Kashyap et al. 2023; Sonkar et al. 2024b).

Municipal wastewater, industrial discharges, and agricultural runoff introduces antibiotics, metals, biocides, and nutrients into rivers, that potentially restructure microbial communities and favoring resistance-associated traits (Martínez 2008; Gao et al. 2025). Non-antibiotic stressors, particularly metals and biocides, may contribute to ARG persistence through co-selection, either *via* co-resistance through physical genetic linkage or cross-resistance through shared physiological mechanisms such as multidrug efflux (Baker-Austin et al. 2006; Seiler and Berendonk 2012). Mobile genetic elements (MGEs), including plasmids, transposons, and integrons, can facilitate the capture, recombination, and horizontal dissemination of resistance determinants (Partridge et al. 2018). Advances in contig-resolved metagenomics have enabled ARGs to be examined within their local genomic contexts, including their physical co-occurrence with biocide and metal resistance genes (BMRGs) and MGE-associated sequences in river sediments (Chen et al. 2019; Gao et al. 2025a). Virulence-factor (VFs) genes may also occur within related genomic regions that combine resistance, mobility, and potential pathogenicity (Zou et al. 2023a). Despite these advances, environmental AMR studies remains dominated by read-based abundance profiles, which cannot establish genetic linkage or distinguish shared environmental exposure from true genomic co-location (Gillieatt and Coleman 2024; Mao et al. 2024). Contig-level analysis can partially address this limitation by identifying ARGs and other determinants on the same assembled sequence (Chen et al. 2019). However, same contig occurrence or proximity alone does not demonstrate co-expression, direct functional interaction, active horizontal transfer, or in-situ selective advantage (Arredondo-Alonso et al. 2017). Therefore, taxonomic, genomic, mobile, and functional relationships among ARGs, BMRGs, MGEs, and VFGs in urban river sediments remain insufficiently resolved.

River sediments retain particle-associated microorganisms, extracellular DNA, and catchment-derived contaminants, potentially integrating deposited inputs over longer timescales than individual water-column samples. Studies on Indian rivers have reported diverse sediment resistomes and clinically relevant ARGs at wastewater-impacted sites (Marathe et al. 2017; Rout et al. 2024, 2026; Sharma et al. 2024), although their sources and drivers vary among river systems. Shotgun metagenomics enables culture-independent characterization of sediment resistome (Marathe et al. 2017; Palit and Das 2020). Read-based analyses quantify ARG abundance and composition, whereas assembly-based approaches recover local genomic neighborhoods and distinguish multiple annotations to a single open reading frame (ORF) from distinct genes cooccurring on the same contig (Grossart et al. 2020). This distinction is important for evaluating associations among ARGs, BMRGs, MGE, and VFs-associated determinants (Chen et al. 2019b).

This study investigated sediment metagenomes from two contrasting wastewater-impacted river systems in India using a staged, provenance-resolved framework. The main objectives were to: (i) characterize ARG diversity while retaining sample-, contig-, and ORF-level provenance; (ii) distinguish same-ORF dual annotations from associations between distinct ORFs on the same contig; (iii) evaluate plasmid-related, MGE, BMRG, VFG, and nitrogen-cycling contexts of ARG-carrying contigs (ACCs); and (iv) classify ACCs into supported taxonomic ranks. This framework explicitly prioritizes ARG loci for investigation based on genomic proximity, mobility-related evidence, and taxonomic resolution.

## 2. Materials and methods

### 2.1 Study area and sediment sampling

Sediment samples were collected from two anthropogenically impacted urban rivers in India the Musi River, Hyderabad, Telangana and the Adyar River, Chennai, Tamil Nadu (**Fig. 1**, **Table S1**). The Musi River, a major tributary of the Krishna River, receives approximately 1.25 million m^3^ day^-1^ of untreated and partially treated municipal wastewater, in addition to industrial effluents (Jampani et al. 2018; Patel et al. 2023). Its catchment forms part of the interior Deccan Plateau within the Krishna River Basin that lies at elevations of approximately 300–600 m (Patel et al. 2023) and exhibits a gradual eastward regional slope (Jogu and Gardas 2025). Four sites distributed along an approximately 140 km downstream reach were selected to represent contrasting anthropogenic settings. Nagole (NH), located immediately downstream of major urban wastewater discharges; Nagaram (NRH) and Musi Reservoir (MRH), representing intermediate reaches; and Shunya Pahad (SPH), a downstream agricultural and aquaculture area with potential human exposure pathways (Sonkar et al. 2024a).

**Figure 1.**
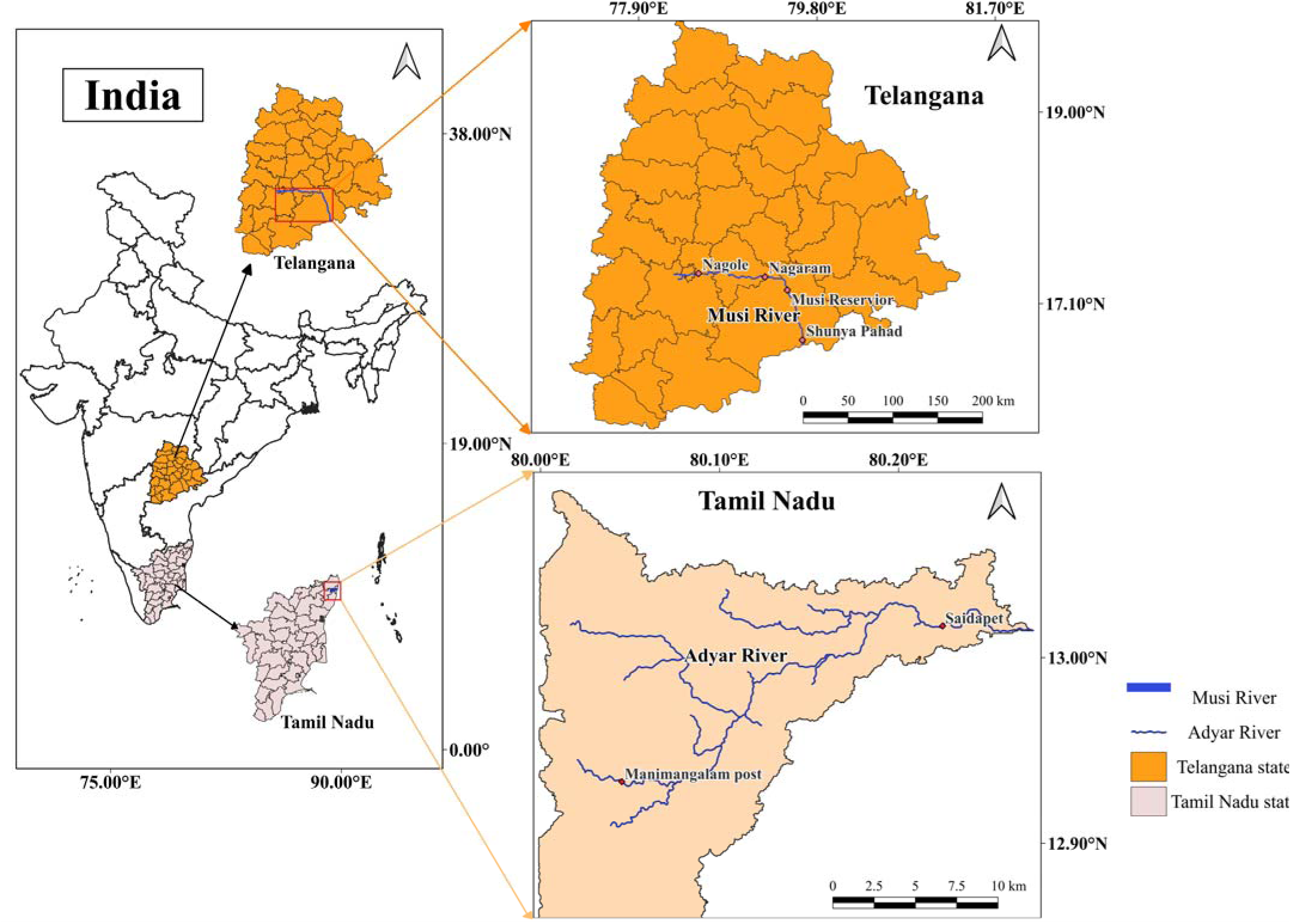
Geographical location of study areas showing sediment sampling points along the Musi River basin in Telangana and the Adyar River basin in Tamil Nadu, India.

In contrast, the Adyar River originates within the upstream tank system near Manimangalam receives surplus flow from Chembarambakkam Lake (12.932° N, 79.997° E), and extends approximately 42 km before discharging into the Bay of Bengal. Its approximately 860 km² catchment is characterized predominantly by flat terrain with a gentle eastward slope (Ramachandran et al. 2019; Saravanan et al. 2019). The river is strongly influenced by municipal sewage and urban runoff, with comparatively low to no industrial pollution loads (Jayaprakash et al. 2012). Two sites were selected: Manimangalam (MM) representing an upstream reach associated with lake-and-tank catchment, and Saidapet (SM) representing a densely urbanized downstream reach hydrologically connected to the estuarine zone. Together, these six sites provided a comparative framework for evaluating sediment-associated AMR under different anthropogenic pressures. Surface sediment samples were collected from a depth of 0–10 cm between February and April 2021 and transferred into pre-autoclaved polypropylene bags. Samples were transported to the laboratory on ice at 4 °C and stored at −20 °C until further analysis. Collectively, these six sites across both rivers represented by contrasting anthropogenic settings and provided a comparative assessment for evaluating sediment-associated AMR and its potential public-health relevance.

### 2.2 Sample processing and shotgun metagenomic sequencing

Total genomic DNA was extracted from 250 mg of each sediment sample using the NucleoSpin® Soil Kit (Macherey-Nagel, Germany) according to the manufacturer’s instructions. DNA quality and concentration were assessed using a NanoDrop spectrophotometer (Thermo Fisher Scientific, USA) and verified by 0.8% agarose gel electrophoresis. Samples with A260/A280 and A260/A230 ratios between 1.8 and 2.1 were considered suitable for library preparation. Paired-end sequencing libraries were constructed from 100 ng of high-quality genomic DNA using the Illumina TruSeq Nano DNA Library Preparation Kit (Illumina, USA). Genomic DNA was fragmented to an average insert size of approximately 350 bp using a Covaris M220 ultrasonicate (Covaris, USA), and library concentrations were quantified using a Qubit fluorometer (Thermo Fisher Scientific, USA). Libraries were sequenced on the Illumina NextSeq 500 platform using a 2 × 150 bp paired-end sequencing strategy, generating approximately 5 Gb of raw sequence data per sample. Library preparation and sequencing were performed by Eurofins Genomics India Pvt. Ltd. (Bengaluru, India).

### 2.3 Metagenomic assembly and gene prediction

Raw paired-end reads were assessed using FastQC (v0.11.9), and quality-control reports were summarized with MultiQC. Adapter trimming, quality filtering, and removal of low-quality or ambiguous bases were performed using fastp (v1.0.1) with default parameters (Chen 2025). High-quality reads from each sample were assembled independently using MEGAHIT (v1.2.9) with default parameters (Li et al. 2016). Assembly quality was evaluated using MetaQUAST (v5.3.0), and total assembly length, N50, and contig counts were recorded for each sample (Mikheenko et al. 2016, 2023). Contigs ≥ 500 bp were retained for downstream analysis.

Open reading frames (ORFs) were predicted using Prodigal (v2.6.3) in metagenomic mode (-p meta) (Hyatt et al. 2010). Predicted protein sequences from all six samples were pooled and clustered using CD-HIT (v4.8.1) at 95% amino acid identity and 90% coverage of the shorter sequence to generate a cross-sample non-redundant (NR) protein catalog (Li and Godzik 2006; Li et al. 2012). The corresponding representative nucleotide ORF sequences were retrieved to construct the mapping reference. Clean reads from each sample were mapped to this catalog using Bowtie2 (v2.5.5) in end-to-end sensitive mode (Langmead and Salzberg 2012). Alignments were processed using SAMtools (v1.23.1), and the resulting mapping statistics were used to estimate ORF coverage (Danecek et al. 2021). Detailed sequencing processing and mapping criteria are provided in **Table S2**.

### 2.4 ARG identification and abundance quantification

Predicted protein sequences were screened against the Structured Antibiotic Resistance Gene database (SARG v3.0) using Diamond (v2.2.5.185; (Buchfink et al. 2014)) (Yin et al. 2023). ARG-like ORFs were identified using thresholds of ≥ 60% amino acid identity, ≥ 70% query coverage, and an e-value ≤ 1 × 10^-10^. Up to five candidate matches were initially retrieved, and the best supported annotation per ORF was retained. An ARG-like ORF was considered reportable in a sample when its representative sequence was supported by at least three mapped reads and ≥ 50% breadth of coverage. Complete search and filtering criteria are provided in **Table S3**. ARG annotations were summarized at the resistance type and subtype levels.

ARG abundance was estimated from the clean reads mapped to the NR-ORF catalog. For each ARG type or subtype, normalized coverage was calculated as:

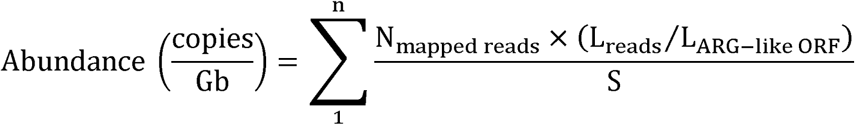

where *N_i_* is the number of clean reads mapped to ARG-like ORF *i*,, *L_reads_* is the mean mapped-read length in base pairs (bp), *L_ARG-like ORF_* is the length of the corresponding ORF (bp), *n* is the number of ORFs assigned to the ARG type or subtype (bp), and *S* is the clean sequencing data volume for that sample in gigabase (Gb). This metric represents sequencing depth normalized ARG coverage and should not be interpreted as an absolute copy number per cell or per gram of sediment.

The potential public-health relevance of the detected ARGs was assessed using the omics based risk-ranking framework of Zhang et al. (2021). Best hit SARG v2.0 accessions were cross-referenced against the SARG risk-ranking resource, and successfully matched ARGs were assigned to Ranks I–IV (Yin et al. 2023). Rank I comprised human-associated, potentially mobile ARGs already detected in pathogenic hosts and therefore represented the highest risk category. Rank II included human-associated and potentially mobile ARGs detected in non-pathogenic hosts, representing potential future threats. Rank III comprised human-associated ARGs without mobility evidence, whereas Rank IV included ARGs lacking evidence of human-associated enrichment. ARG variants without a valid accession-level match or assigned risk rank were retained as unassessed rather than assigned to a lower risk category.

### 2.5 Characterization of ACCs and their genomic neighborhoods

Contigs containing at least one ARG-like ORF were designated ACCs, whereas ACCs containing two or more distinct ARG-like ORF were classified as multi-ARG ACCs. All ORFs represented on ACCs were screened for MGEs (Pärnänen et al. 2018), BMRGs (Pal et al. 2014), VFDB (Zhou et al. 2025), nitrogen-cycling genes (NCycGs; (Tu et al. 2019)). Database specific thresholds and filtering criteria are provided in **Table S3**. In brief, BMRG and VFG annotations required ≥ 70% and ≥ 80% amino acid identity, respectively, with ≥ 70% query coverage and an e-value ≤ 1 × 10^-5^. MGE-associated sequences were identified using BLASTN with ≥ 80% nucleotide identity, an e-value ≤ 1 × 10^-5^, and a minimum alignment length of 75 bp. Primary NCyc annotations required ≥ 85% amino acid identity, ≥ 80% query coverage, and an e-value ≤ 1 × 10^-5^; a relaxed screen ≥ 70% identity, ≥ 70% coverage, and an e-value ≤ 1 × 10^-3^.

For each ARG-MGE, ARG-BMRG, or ARG-VF annotation pair on the same ACC, the genomic relationship was classified as (i) same-ORF dual annotations, (ii) distinct ORFs with overlapping coordinates, (iii) distinct non-overlapping ORFs separated by ≤ 5 kb, or (iv) distinct ORFs separated by > 5 kb. Same ORF dual annotations were distinguished from associations between separate ORFs because they do not represent physically distinct genes. NCycGs were retained as broader functional context annotations and assessed separately.

All retained assembled contigs were screened using geNomad (v1.12.0; database v1.9) end-to-end workflow (Camargo et al. 2023). Contigs classified by geNomad as plasmid-associated were designated plasmid-like. This sequence based designation does not demonstrate the presence of a complete, viable, or conjugatively transferable plasmid.

### 2.6 Taxonomic annotations of ACCs

Putative taxonomic affiliations of ACCs were assigned using the Contig Annotation Tool (CAT) against the CAT nr reference database (dated on 12 December 2024; https://tbb.bio.uu.nl/tina/CAT_pack_prepare/) (Von Meijenfeldt et al. 2019) using the default parameters (**Table S4**). CAT integrates protein level taxonomic evidence across multiple ORFs on each contig and assigns the contig to the lowest taxonomic rank supported by the combined evidence. Assignments were therefore reported at the species, genus, family, order, class, or phylum level, with unsupported assignments retained as unresolved. Species- and genus-level classifications were interpreted as putative contig affiliations rather than confirmed ARG-host identities because short and fragmented contigs, shared genes, and incomplete reference representation can limit taxonomic specificity. Unresolved ACCs were retained in all summaries to avoid classification related reporting bias. Since CAT was applied only to selected ACCs rather than complete metagenomic assemblies, the resulting profiles were used to describe resistome-associated taxonomic contexts and not whole community composition.

Unique, provenance-resolved ACCs were screened for potential human pathogenic bacteria (HPB) by comparing CAT-derived assignments with the HPB species reference list compiled by Zhou et al. (2019). Exact, standardized correspondence between the CAT species assignment and an HPB-listed species was classified as a species-level match, whereas correspondence restricted to the genus-level was treated as suggestive. Candidate mobile-context evidence was defined as either: (i) a geNomad plasmid-like, virus-like, or provirus-region prediction; or (ii) distinct ARG-like and MGE-like ORFs located within 5 kb on the same ACC.

ACCs were assigned hierarchically to mutually exclusive categories in the following order: C1, species-level HPB match with mobile-context evidence; C2, species-level HPB match without mobile-context evidence; C3a, genus-level HPB correspondence with mobile-context evidence; C3, mobile-context evidence without HPB correspondence; C4, genus-level HPB correspondence without mobile-context evidence; and C5, neither HPB correspondence nor mobile-context evidence. These categories summarize sequence-based evidence for taxonomic correspondence and candidate mobility context. They do not confirm pathogenicity, ARG expression, horizontal transfer, host viability, or infection risk.

### 2.7 Data analysis

Data processing, statistical analyses, and visualization were performed in R (v4.5.1) using iNEXT (v3.0.2), vegan (v2.7.2), compositions (v2.0.9), zCompositions (v1.6.0), ggplot2 (v4.0.3), and pheatmap (v1.0.13). ARG subtype richness and diversity were calculated from reportable mapped read counts using Hill numbers of orders (q = 0, 1, 2), corresponding to observed subtype richness, Shannon-effective diversity, and inverse Simpson-effective diversity, respectively. Sampling completeness and coverage based standardized diversity were evaluated using iNEXT (v3.0.2) at the highest common sample coverage supported across samples (Hsieh et al. 2016).

Compositional differences among sediment ARG-subtype profiles were characterized using Aitchison distances calculated as Euclidean distances after centered log-ratio (CLR) transformation (Gloor et al. 2017; Quinn et al. 2018). Before transformation, zero entries were replaced with a single constant equal to half the smallest positive abundance across the analyzed subtype abundance matrix (Quinn et al. 2019). Principal coordinate analysis and average-linkage hierarchical clustering of the Aitchison distance matrix were used to visualize site-associated patterns (Clarke 1993). PERMANOVA was restricted to an exploratory sensitivity analysis, enumerating all 15 distinct allocations of river labels while preserving the two versus four group sizes (Anderson 2001). Separately, SIMPER based on Bray-Curtis dissimilarity was used descriptively to identify subtype contributions (CLARKE 1993). No differential abundance tests, taxon-ARG networks, or confirmatory river level models were fitted. All patterns were interpreted as exploratory.

## 3. Results and discussion

### 3.1 Sequencing output, assembly and read-mapping statistics

Across the six sediment metagenomes, 274,519,498 quality-filtered reads, representing 40.775 Gb of clean sequence data, were retained for analysis (**Table 1**). Sample-level sequence yields ranged from 4.409 Gb at SM to 8.302 Gb at SPH, with mean post-filtering read lengths of 147.85-148.94 bp. After restricting the assemblies to contigs ≥ 500 bp, 1,504,313 contigs were retained, representing a total assembled length of 1.492 Gb. The number of retained contigs ranged from 58,988 at MM to 409,632 at NRH, while assembly N50 ranged from 817 bp at SPH to 1,620 bp at MM. The longest contig was recovered from MM and measured 240,391 bp. Overall, 2,530,998 ORFs were predicted from the retained contigs.

**Table 1.**
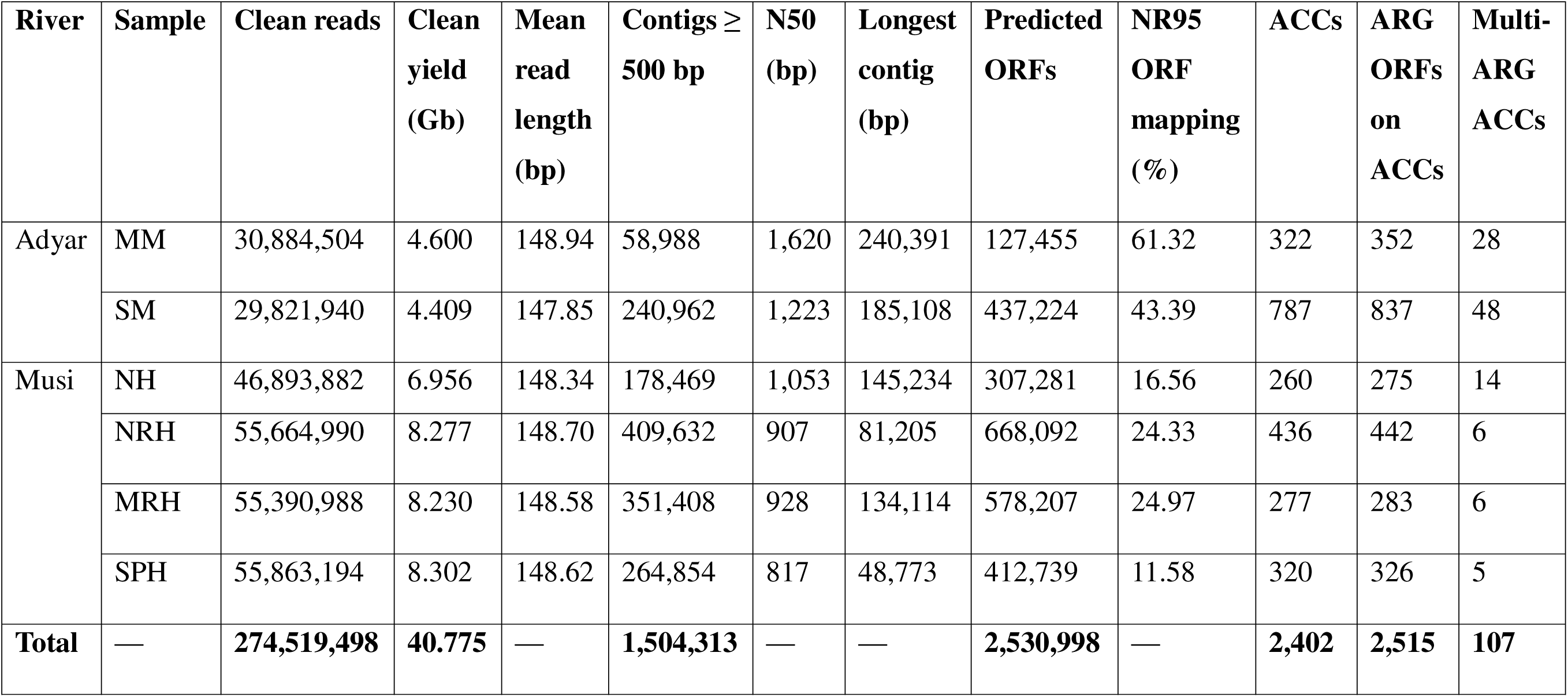
Sequencing and assembly statistics of all sediment metagenomes.

The two Adyar metagenomes comprised 60,706,444 clean reads, corresponding to 9.009 Gb of sequence data and 299,950 retained contigs. MM contained 58,988 contigs and 322 ACCs, whereas SM contributed 240,962 contigs and 787 ACCs. Clean read mapping rates to the NR-ORF catalog were 61.32% at MM and 43.39% at SM.

The four Musi metagenomes comprised 213,813,054 clean reads, 31.766 Gb of sequence data and 1,204,363 retained contigs. NH, NRH, MRH, and SPH contained 178,469, 409,632, 351,408, and 264,854 contigs, of which 260, 436, 277, and 320 were ACCs, respectively. Clean-read mapping rates to the NR-ORF catalog ranged from 11.58% at SPH to 24.97% at MRH.

Overall, read-mapping rates were consistently higher in the Adyar metagenomes than in the Musi metagenomes, indicating that a greater proportion of Adyar reads was represented in the cross-sample NR-ORF catalog. Notably, the downstream Musi site, SPH, had both the lowest mapping rate and the lowest assembly N50 value (**Table 1**), despite generating the largest clean-sequence yield (8.302 Gb). These contrasting metrics indicate that sequencing yield alone did not determine assembly contiguity or catalog representation.

### 3.2 ARG occurrence and diversity

Applying reportable detection filtering (≥ 1 ORF supported by ≥ 3 mapped reads spanning ≥ 50% length) retained 90.71–97.96 % of normalized ARG abundance across all six sediment metagenomes, confirming that low support annotations contributed negligibly to overall abundance estimates (**Fig. S1A**). Overall, 264 of 280 evaluated ARG subtypes were reportably detected in at least one sample, with 18 core subtypes ubiquitous across all sites and 92 classified as sample-exclusive (**Fig. S1B**). Between basins, 138 subtypes were shared across both river systems, whereas 75 and 51 subtypes were exclusive to Adyar and Musi sediments, respectively (**Fig. S1B**).

Adyar River sediments contained richer and more abundant resistomes (MM: 124 subtypes, 1,010.09 ×/Gb; SM: 171 subtypes, 1,295.90 ×/Gb) than Musi River sediments (130.47–225.67 ×/Gb), where subtype richness decayed progressively downstream (NH: 110; NRH: 92; MRH: 67; SPH: 65) (**Fig. 1A**, **Fig. 4A**). Elevated ARG abundance and richness in urban-impacted sediments are consistent with Marathe et al. (2017), which documented enrichment of ARGs, MGEs and potential pathogens in sewage-impacted Mutha River sediments. However, environmental ARG enrichment can frequently reflect the introduction of sewage-associated microbial biomass rather than local selection alone (Berendonk et al. 2015; Karkman et al. 2018), and the relative contributions of incoming biomass, resident communities, and contaminant exposure cannot be separated here.

Multidrug efflux annotations had the highest mean normalized abundance across all samples (195.83 ×/Gb), followed by aminoglycoside resistance (110.37 ×/Gb) (**Fig. 1B**). However, dominant subtypes shifted markedly by location (**Fig. 2, Fig. S1C**). The efflux gene *mexB* led at SM (75.37 ×/Gb) and NH (12.64 ×/Gb), whereas *aph(3′)-IIIa* dominated at MM (198.96 ×/Gb), identifying aminoglycoside phosphotransferase-mediated resistance as a major component of its profile (Zhang et al. 2023). Downstream Musi sites transitioned to *bacA* at NRH (21.45 ×/Gb) and *novA* at MRH (13.43 ×/Gb) and SPH (18.67 ×/Gb), indicating changes in the composition of resistance-associated functions. Crucially, the predominance of these baseline determinants does not establish indigenous host origin, lower clinical relevance, or disappearance of wastewater influence.

**Figure 2.**
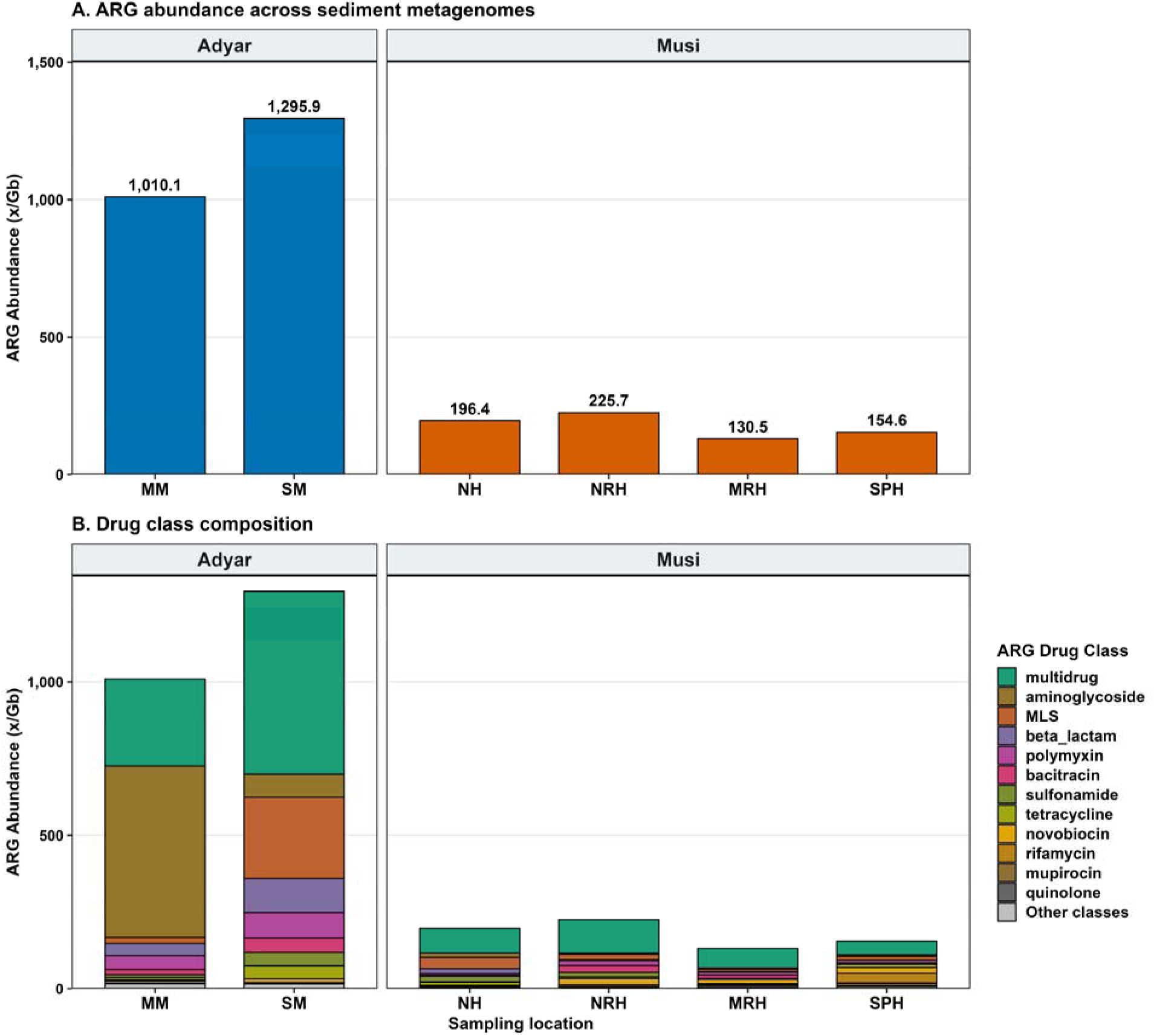
**(A)** Total reportable ARG abundance, and **(B)** Drug class composition across the sediment metagenomes.

**Figure 3.**
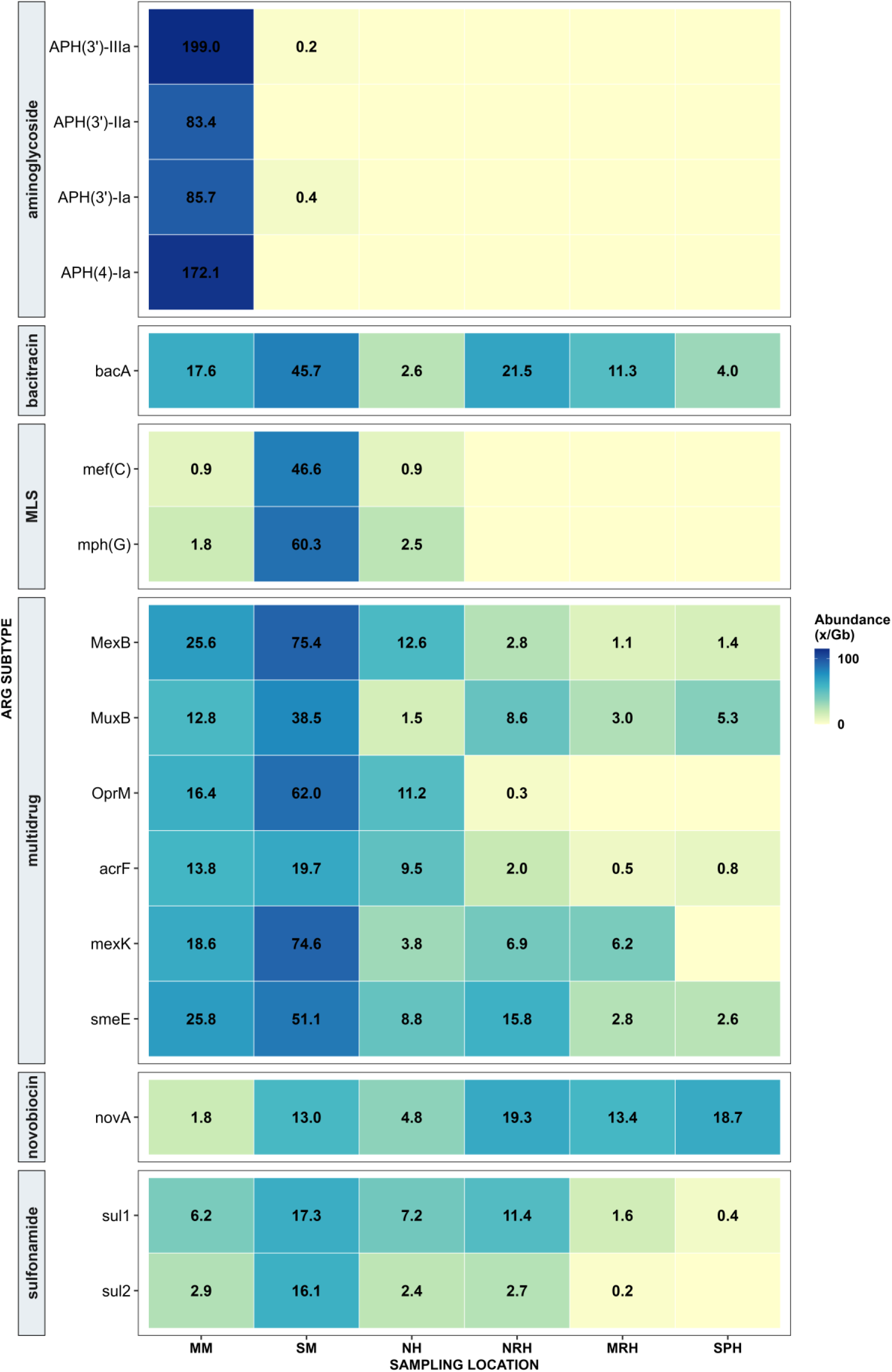
Heatmap of leading ARG subtypes grouped by drug class across sediment samples.

Hill-number profiles (**Fig. 4A, Fig. S2A and S2B**) distinguished subtype richness from abundance-weighted diversity, expressed as effective numbers of equally abundant subtypes (Jost 2006; Ott et al. 2021). Adyar downstream site (SM) had the highest richness, Shannon- and Inverse Simpson-effective diversity (171, 68.78 and 41.20), respectively. Although Musi upstream (NH) contained fewer subtypes than Adyar upstream (MM) (q_0_ = 110 vs. 124), it retained substantially higher effective diversity (q_1_ = 60.60 vs. 28.32; q_2_ = 40.06 vs. 11.56). This contrast demonstrates that MM profile was more strongly concentrated in a limited number of ARG subtypes, whereas NH supported a more balanced abundance distribution. Unweighted richness alone therefore understated the structural differences between these resistomes.

**Figure 4.**
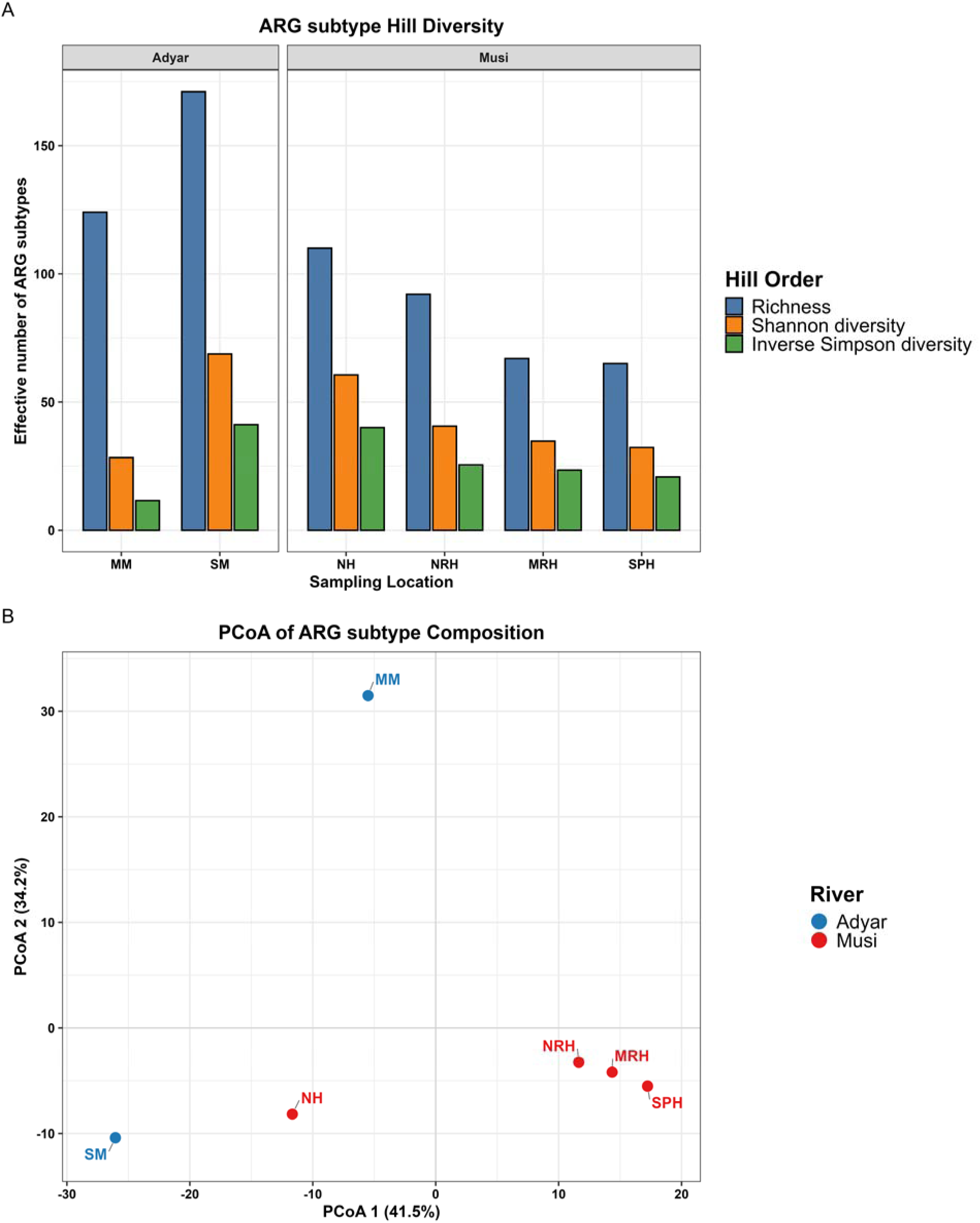
**(A)** ARG subtype diversity and **(B)** PCoA of ARG subtype composition at each sampling location.

CLR Aitchison ordination further captured spatial heterogeneity, with the first two PCoA axes explaining 75.74% of resistome variance (PCoA1: 41.5%; PCoA2: 34.2%; **Fig. 4B**). Site MM separated distinctly along PCoA2, consistent with its uneven profile and prominent aminoglycoside component. Pairwise Aitchison distances identified the downstream Musi samples as the most compositionally similar and both Adyar samples as the most dissimilar (**Fig. S2C and S2D**). Separately, Bray-Curtis SIMPER decomposition identified aminoglycoside (*aph(3’)-IIIa*, *aph(4)-Ia*), novobiocin (*novA*), multidrug (*MexF*), bacitracin (*bacA*), and other multidrug-efflux annotations as major contributors to dissimilarity rather than river-specific markers (**Fig. S2E**). While the lower richness, reduced abundance, and compositional clustering downstream in the Musi align with reduced upstream inputs, microbial turnover, or hydrological dilution (Gweon et al. 2021; Lee et al. 2021). It should not, however, be described as attenuation through dilution or deposition because each location was sampled once and no discharge, travel time, or longitudinal mass-balance data were evaluated.

### 3.3 ARG risk-ranking profiles

Across the six sediment metagenomes, cumulative reportable resistome abundance was 3,013.07 ×/Gb (**Table S5**). Low-risk Rank IV determinants accounted for 2,344.88 ×/Gb (77.82%), followed by high-risk Rank I (298.02; 9.89%), Rank II (199.88; 6.63%), Rank III (140.80; 4.67%) and unassessed variants (29.51; 0.98%) (**Fig. 5A, B; Table S5**). Full annotation completeness metrics confirmed complete coverage of all 2,055 non-redundant ARG representatives against the reference database (**Table S6**). This predominance of low-risk determinants highlights why raw ARG abundance alone is insufficient to establish surveillance priorities. Nevertheless, these rank classification does not establish that the detected determinants are intrinsically harmless, functionally inactive or permanently non-mobilizable. Zhang et al. (2021) explicitly identified limitations of the enrichment criterion, including the classification of few clinically important sequences as Rank IV because of their widespread environmental occurrence. Conversely, Rank I assignment does not demonstrate carriage by a viable pathogen or presence on a mobile element in these samples. Risk ranks therefore complement, rather than replace, sample-specific genomic-context and host analyses (Martínez et al. 2015; Zhang et al. 2021).

**Figure 5.**
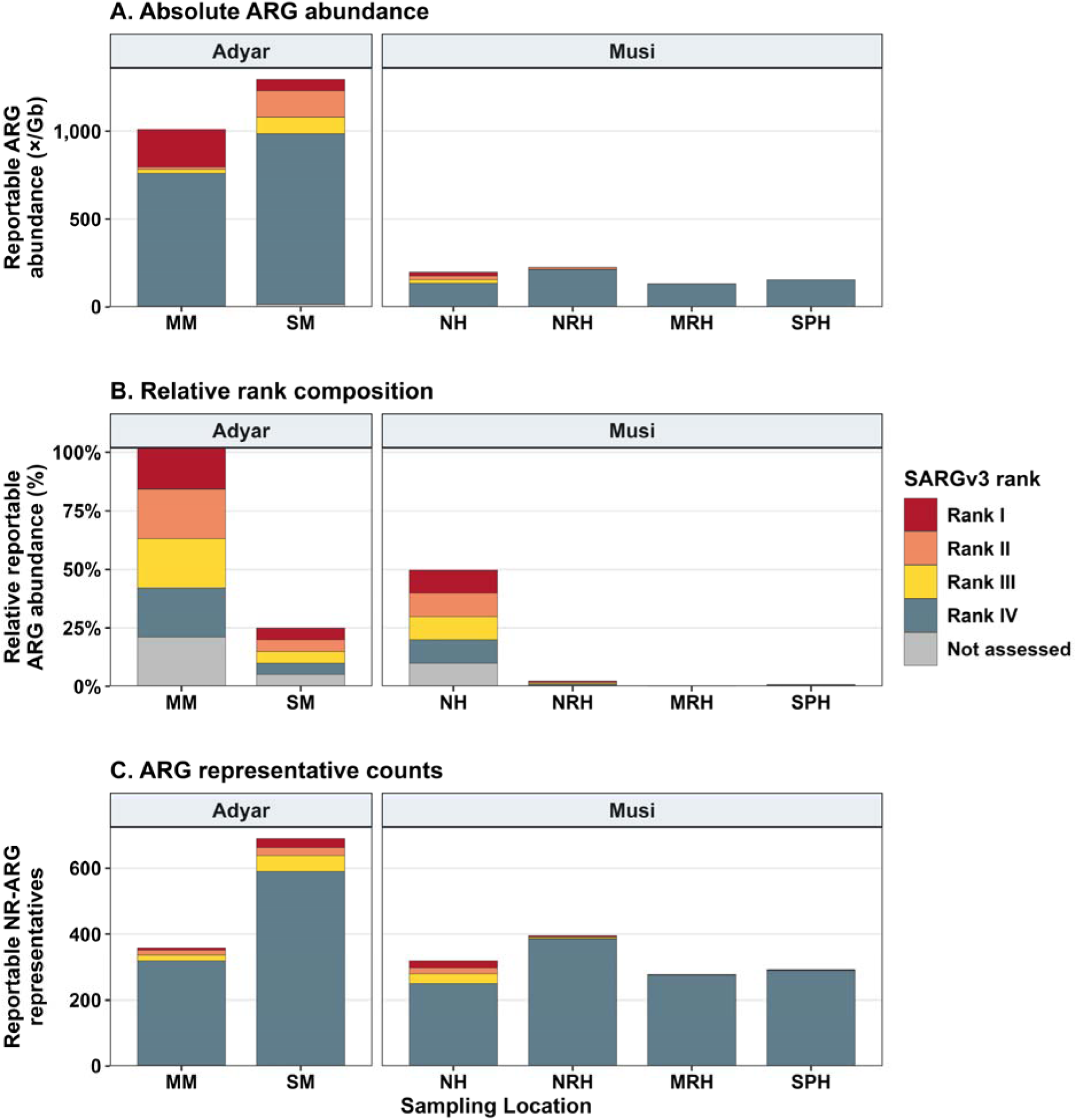
SARG v3 based risk-rank profiles of ARG-like ORFs across sediment metagenomes. **(A)** Absolute reportable ARG abundance; **(B)** Relative rank composition of ARG abundance; **(C)** Number of reportable NR-ARG representatives assigned to each rank.

Rank IV determinants remained the dominant abundance category at every site, ranging from 128.55 at MRH to 969.59 ×/Gb at SM (**Fig. 5A; Table S5**). Spatially, Rank I determinants were heavily concentrated, rather than exclusively restricted to urban-impacted reaches. High-risk Rank I abundance was highest at upstream Adyar site MM (212.63), followed by its downstream site SM (64.53) and Musi urban site NH (19.54). Much lower abundances occurred at NRH (1.06) and SPH (0.26), with no reportable detection at MRH (**Fig. 5A, B**). Rank II and III abundances were greatest at SM (151.50 and 93.13, respectively), followed by NH (21.55 and 22.36, respectively). Reported Rank II abundances were low (0.40–1.43), and Rank III representatives were not reportably detected at downstream Musi (SPH). Unassessed abundance ranged from 0 at SPH to 17.16 ×/Gb at SM, representing these variants remain unclassified rather than demonstrably low-risk.

Richness profiling of reportable NR-ARG sequences provided complementary structural insights (**Fig. 5C; Table S7**). Site SM contained the broadest representation across all risk tiers (27 Rank I, 25 Rank II, 48 Rank III, and 588 Rank IV sequence representatives), followed by NH (22, 17, 30, and 248 representatives, respectively). These counts represent sequence-level richness rather than ARG subtype richness (**Fig. S3**). Comparing risk-tier richness with abundance-weighted diversity highlights the necessity of multi-dimensional resistome auditing. SM therefore combined broad tier representation with high overall diversity, whereas MM contained the greatest Rank I abundance despite its less even resistome (**Fig. 5A–C; Table S5 and S7**). This contrast shows why surveillance priority should consider the abundance and identity of high-risk determinants rather than total ARG abundance or richness alone (Lee et al. 2022).

The lower Rank I signals at Musi downstream indicate reduced detectable representation rather than attenuation or elimination from the aquatic system. Differences in inputs, hydrological dilution and retention, sediment redistribution, microbial community turnover, assembly recovery limitations, and database coverage may influence detection (Chen et al. 2020; Lee et al. 2022). The spatial profiles therefore identify urban-associated sites for targeted genomic and exposure assessment without demonstrating downstream removal or sediment to host transmission (Gweon et al. 2021; Jiang et al. 2024).

### 3.4 Mobility-associated genomic context of ACCs

Among 2,402 evaluated ACCs, 75 (3.12%) harbored at least one qualifying mobile-context signal, defined as a geNomad-predicted plasmid-like or virus-like classification, a predicted proviral region, or an ARG-like ORF situated within 5 kb of a distinct MGE-annotated ORF. These non-mutually exclusive categories comprised 53 plasmid-like ACCs, six virus-like ACCs, two ACCs containing provirus regions, and 36 ACCs containing proximal ARG-MGE-reference pairs (≤ 5 kb) (**Table S8; Fig. S4**). These signals reflect mobility-associated genomic features rather than horizontal transfer frequencies. Moreover, short-read assembly fragmentation limits recovery of neighboring genetic elements, so ACCs lacking detectable mobility signatures cannot be assumed to be strictly chromosomal or non-transferable (Brown et al. 2022).

Spatially, absolute counts and proportional representations revealed complementary aspects of mobilome architecture. For the Adyar, site SM yielded the largest absolute count of context-positive ACCs (32/787; 4.07%), whereas MM exhibited the highest proportional representation (18/322; 5.59%). Within the Musi River, the proportion of ACCs carrying mobility-associated evidence decreased descriptively from NH (14/260; 5.38%) to NRH (10/436; 2.29%) and MRH (1/277; 0.36%), with zero mobile-context signals at SPH (0/320) (**Table S8; Fig. S4**). This contrast demonstrates that although SM contributed the largest absolute number of context positive ACCs, MM and NH showed the highest proportional representation of mobility-associated contexts within their respective ACC inventories.

Across all ACCs, 48 ORFs received MGE-reference annotations, comprising 36 insertion-sequence/transposition-associated enzymes, eight qacEΔ-like elements (representing integron-associated resistance markers rather than mobility machinery), and four integrase-like ORFs (**Table S9**). The *qacEΔ*-like annotations were treated as integron-associated resistance markers rather than independent mobility machinery. Coordinate analysis identified 54 distinct-ORF ARG-MGE pairs, of which 52 occurred within 5 kb across 36 ACCs (**Table S10**). Ten specific ARG subtype-MGE category pairings recurred across at least two ACCs (**Fig. 6A; Table S11**). The most frequent co-localization linked *sul2* to transposition-associated features on five ACCs spanning SM (1), MM (1), NH (1), and NRH (2). Aminoglycoside resistance determinants *aph(3″)-Ib* and *aph(6)-Id* were each physically linked to transposition features across three ACCs spanning SM, MM, and NH. Other recurrent co-localizations involved *sul1* across Adyar ACCs, *sul4* on two ACCs from SM and NRH, and "other class A *β-lactamase*", *ereA*, and *linG*, each on SM and NH (**Fig. 6B; Table S11 and S12**). Recurrence across geographically separated rivers aligns with established resistance associated with mobile element architecture (Partridge et al. 2018).

**Figure 6.**
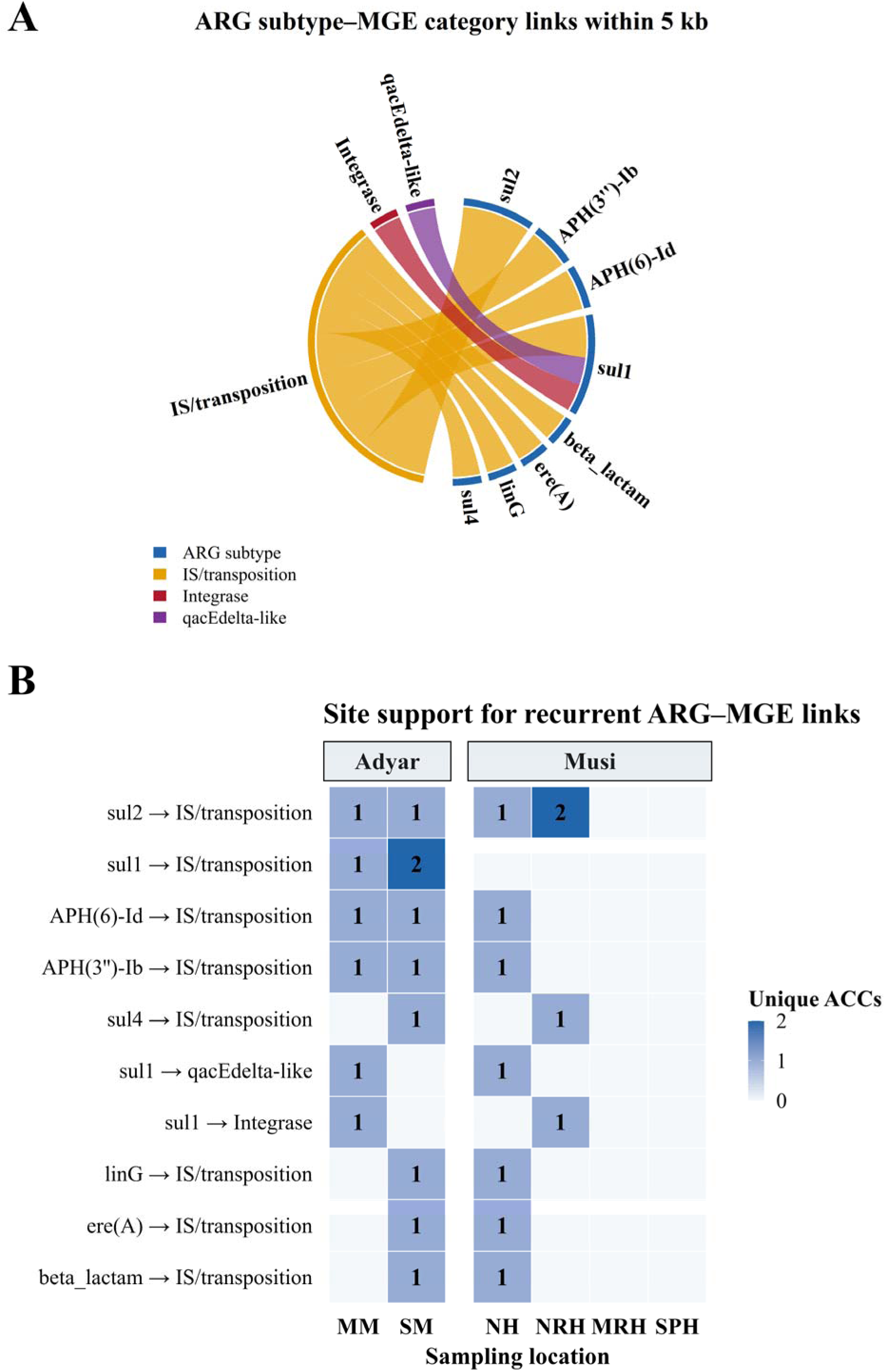
**(A)** Genomic colocalization network of ARG subtypes and MGE categories within a 5 kb genetic window; (B) Recurrent ARG-MGE relationships shared between the Adyar and Musi sediment metagenomes.

Separately, *sul1* occurred near integrase-like annotations on MM and NRH sites and near *qacEΔ1*-like annotations on MM and NH (**Fig. 6B; Table S12**). These pairwise associations are compatible with class 1 integron-associated regions, whose conserved components frequently include *intI1*, *qacEΔ1*, and *sul1* (Gillings et al. 2015). However, because these annotations were recovered on fragmented short-read contigs, they do not establish recovery of a complete canonical *intI1*-*qacEΔ1-sul1* region. Although clinical *intI1* abundance is widely recognized as a proxy for anthropogenic pollution and resistome enrichment (Gillings et al. 2015; Czekalski et al. 2015), detecting uncharacterized integrase annotations or partial gene linkages on contigs cannot independently establish that proxy without more complete sequence resolution (Zhang et al. 2018; Chen et al. 2019; Mao et al. 2024).

No ARG-like ORFs overlapped predicted provirus-region boundaries (**Fig. S4; Table S8**). This positional result is distinct from the virus-like sequence classification assigned to six ACCs, which does not independently validate ARG carriage within functional viral genomes. geNomad sequence classification and provirus predictors evaluate different genomic signatures, while permissive ARG annotation or bacterial contamination can inflate apparent phage-associated resistance (Enault et al. 2017). Overall, these recovered co-localizations identify resistance-associated genomic regions with candidate mobility relevance. Establishing functional mobility will require long-read or hybrid sequencing, host-linkage validation, and transfer assays appropriate to the recovered elements; short-range proximity alone does not demonstrate expression, complete transfer machinery, or horizontal gene transfer (Arango-Argoty et al. 2018; Mao et al. 2024).

### 3.5 Genomic co-occurrence and functional context of ACCs

#### 3.5.1 BMRG-associated genomic context

BacMet annotations were identified on 386 of 2,402 canonical ACCs across all six sediment metagenomes. Prevalence was highest at the upstream Adyar site MM (113/322 ACCs; 35.09%), whereas the downstream Adyar site SM contributed the largest absolute count (148/787; 18.81%). Along the Musi River, prevalence decreased from NRH (51 of 436 ACCs; 11.70%), NH (27 of 260 ACCs; 10.38%), through MRH (28 of 277 ACCs; 10.11%) to SPH (19 of 320 ACCs; 5.94%) (**Table S13; Fig. 7A and S5A**). Annotated ACCs were assigned to biocide (356 ACCs), metal (23 ACCs) and combined biocide-metal (16 ACCs) resistance groups, as these counts sum to more than 386 because some ACCs can carry annotations from more than one group. Biocide-associated categories was dominated by acridine (194 ACCs), phenolic compounds (178), phenanthridine (167), organosulfates (150), followed by xanthene (131) and quaternary ammonium compounds (QACs; 127). Metal-associated annotations included zinc (21 ACCs across all sites), mercury (9; MM and NRH), copper (7; SM, MM and NH), chromium (5; SM and MM), and cobalt and nickel (2 each at SM and NH) with single ACCs carrying antimony and arsenic (MM) or iron (NH) (**Table S13**). These reference-associated compound categories describe putative resistance functions.

**Figure 7.**
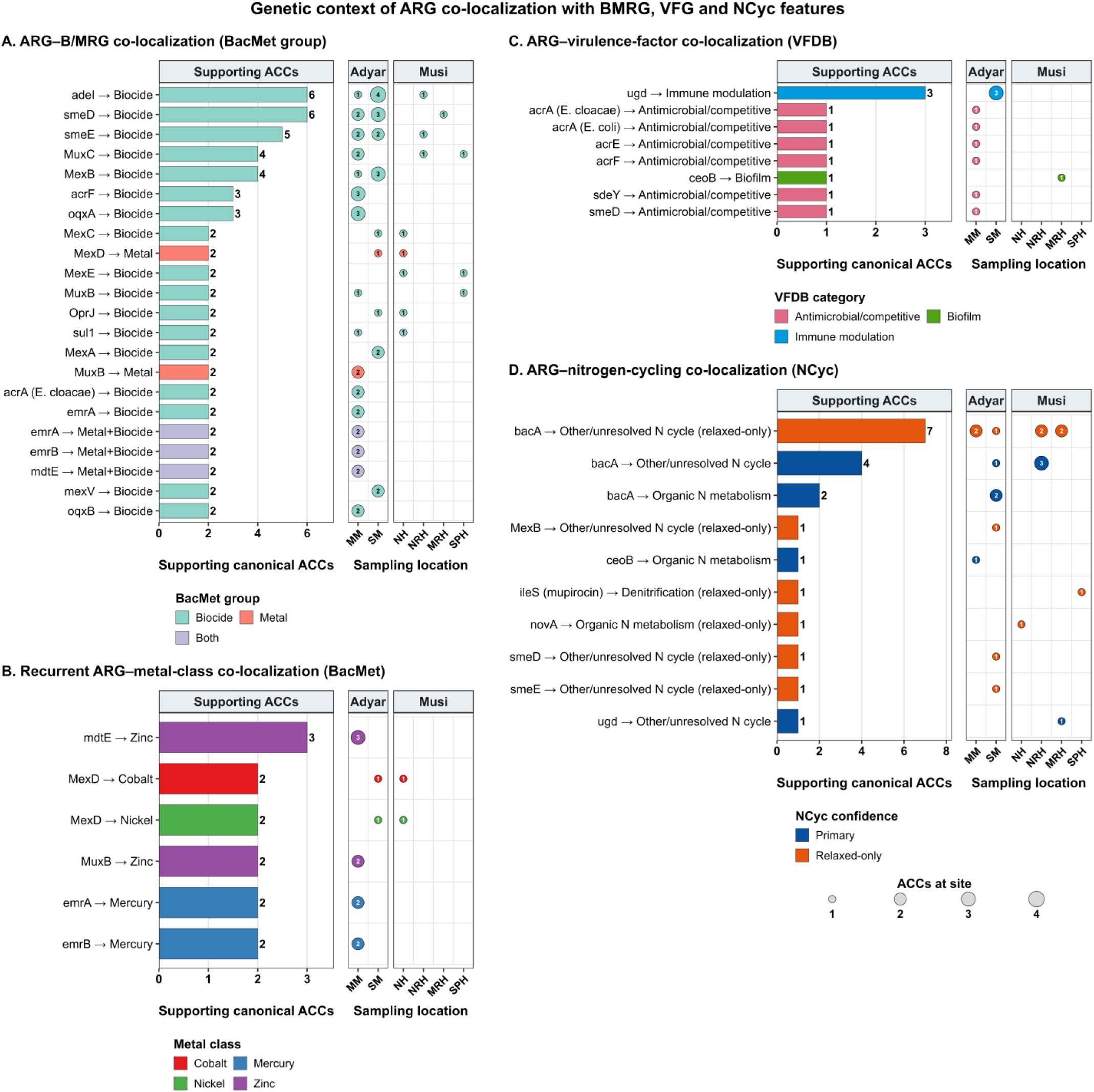
Genomic co-localization of ARGs with **(A)** BMRGs; **(B)** metals class only; **(C)** VFGs; and **(D)** nitrogen cycling genes on assembled contigs across Adyar and Musi river sediments.

Distinguishing single-ORF annotations from neighboring genes substantially narrowed the evidence for ARG-BMRG co-selection (Pal et al. 2015; Li et al. 2017). Of 490 ARG-BMRG annotation pairs, 399 assigned SARG and BacMet labels to the same ORF and were excluded from physical co-localization analysis. These same-ORF overlaps predominantly encode broad-substrate efflux pumps (e.g., RND-family transporters such as *adeI*, *adeJ*, *smeE*, and *MuxC*) and membrane-stress regulators that confer multi-compound physiological tolerance rather than physically linked, independently resistance cassettes (Nikaido 2009; Seiler and Berendonk 2012). Evidence for co-localization of distinct genes was therefore restricted to the remaining 91 ARG-BMRG pairs on different ORFs, all within 5 kb, across 61 ACCs (**Table S14**). Of these pairs, 77 involved separate ORFs and 14 involved adjacent ORFs with overlapping intervals and the latter may include fragments of a single gene and are interpreted cautiously. Their prevalence was highest at MM (31/322; 9.63%) followed by SM (20/787; 2.54%), NH and NRH (3 each), and MRH and SPH (2 each) (**Fig. S5A**). Of the 61 ACCs with a distinct-ORF ARG-BMRG pair, 55 (90.2%) involved multidrug-class ARGs, predominantly neighboring components of multicomponent efflux systems, whereas six ACCs (9.8%) paired non-multidrug ARGs with *qacEΔ1* (*sul1* at MM and NH; OXA-4 at SM), *qacF* (SM), *merR* (two aminoglycoside resistance genes on one MM contig) or *sugE* (MM) (**Table S14**). Restricting the analysis to distinct BMRG ORFs narrowed biocide-class associations to acridine (32 ACCs), organosulfates (30), QACs (23), phenanthridine (25), xanthene (24) and phenolics (18) (**Table S15A**). Metal-class associations at MM comprised zinc (5 ACCs) and mercury (3 ACCs), with single ACCs carrying copper, antimony or arsenic. Alongside chromium occurred on two ACCs spanning both Adyar sites, and cobalt and nickel annotations were each supported by two ACCs at SM and NH, both linked to MexD (**Table S15B; Fig. 7B**).

Notably, a distinct-ORF association between *sul1* and the quaternary ammonium resistance gene (*qacEΔ1*) was detected on two ACCs, one each from MM and NH (**Table S14 and S15C**). In both ACCs the ORF intervals overlap, so this association rests on adjacent, overlapping ORFs. The arrangement is consistent with the 3′-conserved segment of class 1 integron architecture, where *qacEΔ1* and *sul1* are typically adjacent, and *qacEΔ1* and mercury resistance operons are among the few biocide/metal determinants frequently reported on ARGs on mobile elements (Gillings et al. 2015; Pal et al. 2015). However, the present linkages do not independently establish complete integrons, plasmid carriage or active co-selection (Di Cesare et al. 2016). Confirming functional cross-resistance or exposure driven co-selection in sediments requires matched environmental geochemistry (Seiler and Berendonk 2012; Gupta et al. 2022), metatranscriptomics profiling to demonstrate active gene expression (Liu et al. 2019; Zhang et al. 2025a), and culture-based phenotypic validation (Marathe et al. 2017; Karkman et al. 2018).

#### 3.5.2 VF-associated genomic context

VFDB matches occurred on 28 ORFs across 26 ACCs, concentrated at the Adyar sites MM (17/322 ACCs; 5.28%; 19 ORFs) and SM (7/787; 0.89%; 7 ORFs), with minor occurrence at Musi site MRH (2/277; 0.72%) and none across all other Musi sites (NH, NRH, and SPH) (**Fig. 7C**; **Table S16A, B**). At MM, 15 of 19 ORFs were assigned to antimicrobial activity/competitive advantage (*acrB*, 11; *acrA*, 4) and four to immune modulation (*ugd*, 3; *msbA*, 1). At SM, five of seven ORFs were immune modulation (*ugd*, 4; *galU*, 1) and two were biofilm-associated (*adeG*), as were both MRH matches (*adeG*) (**Table S16C**). Of the 28 VFDB-matched ORFs, 27 (in 25 ACCs) were also annotated as ARGs on the same ORF, and only one (*galU*, SM) was a distinct ORF (**Table S16B,D**). The VFDB signal therefore largely reflects efflux and lipopolysaccharide modification genes that carry both ARG and VF annotations, not independent virulence loci co-localized with ARGs. The localized spatial enrichment of VFGs at these urban-impacted Adyar sites is consistent with enrichment of efflux-associated genes reported across heavily disturbed Asian river catchments (Chen et al. 2019; Zhang et al. 2022; Mao et al. 2025).

Matching genes to VFDB requires careful interpretation, because many matches are genes that support bacterial survival in the environment rather than cause human disease (Zou et al. 2023b). Here, VFDB hits were dominated by efflux components (*acrA*, *acrB*, *adeG*) and lipopolysaccharide modification genes (*ugd*, *galU*, *msbA*), which have general roles in envelope integrity, stress tolerance and competition (Liu et al. 2019; Xi et al. 2024). Because 27 of 28 VFDB-matched ORFs were also annotated as ARGs on the same ORF, the overlap reflects shared function between the ARG and VFDB databases rather than physical clustering of independent virulence determinants with ARGs. The concentration of matches at the urban-impacted Adyar sites is consistent with wastewater associated changes in the river resistome and mobilome (Lekunberri et al. 2018; Zhang et al. 2025b), but sequence similarity alone cannot establish pathogenicity, host range or selection, and ARG enrichment in impacted environments may reflect fecal input rather than on-site selection (Karkman et al. 2019).

#### 3.5.3 Nitrogen-cycling functional context

Primary-threshold NCyc annotations were identified on 12 distinct ORFs across 12 ACCs, located at MM (5/322 ACCs; 1.55%), NRH (3/436; 0.69%), SM (3/787; 0.38%), and MRH (1/277; 0.36%), with none detected at NH or SPH (**Fig. 7D; Table S17A**). All 14 ARG-primary NCyc annotation pairs involved distinct ORFs, eight of these were physically co-localized within 5 kb across eight ACCs. The remaining six lay farther apart on the same contig (**Table S17D**). Pathway-resolved annotations comprised organic nitrogen metabolism on two ACCs at each Adyar site (MM: *gdh* and *glnA*; SM: *glnA* on both) and assimilatory nitrate reduction of one ACC at MM (*nasA*) (**Table S17B,C**). The remaining seven ORFs, distributed across MM (2), SM (1), NRH (3) and MRH (1), had no specific nitrogen-cycling pathway assignments and were excluded from pathway level interpretation. Four were glutamate synthase subunit genes at SM (1) and NRH (3) and three carried only a database identifier (MM, 2; MRH, 1) (**Table S17C**).

Relaxed-threshold screening recovered an additional 16 distinct ORFs across 16 ACCs, none shared with the primary set and spanning all six locations. At MM (2/322; 0.62%), SM (5/787 ACCs; 0.64%), NH (1/260; 0.38%), NRH (3/436; 0.69%), MRH (4/277; 1.44%), and SPH (1/320; 0.31%) (**Fig S5C; Table S17A**). Among 17 ARG-relaxed NCyc pairs, four represented same-ORF dual annotations and 13 involved distinct ORFs, 12 of which physically co-localized within 5 kb across 11 ACCs (**Table S17D**). Pathway assignment was limited, as one ORF was annotated as organic nitrogen metabolism (*gdh* at NH), one carried a denitrification labelled gene (*narB* at SPH), and the remaining 14 were unassigned, comprising glutamate synthase subunit hits (6), *hcp* (1 at MRH) and ORFs identified only by database identifier (7) (**Table S17B,C**). No nitrification context was recovered, and a single denitrification labelled ORF was insufficient to infer a complete denitrification pathway in any ACC.

ACCs contained NCyc-annotated ORFs in only eight of 14 primary and 12 of 13 distinct relaxed ARG-NCyc pairs, and pathway resolved annotations were sparse in the primary set, with no nitrification and denitrification pathway in either set. Physical proximity of this kind does not demonstrate metabolic or regulatory coupling between antibiotic resistance and nitrogen transformation. Microcosm experiments have shown that antibiotic exposure can alter nitrate reduction and denitrification, and that nitrate-reducing bacteria host a large share of expressed ARGs (Deng et al. 2026). These findings indicate an ecological association rather than a gene level functional link. Likewise, reported co-hosting of ARGs and denitrification genes (Wang et al. 2024; Deng et al. 2026) concerns dissimilatory nitrate-reducing taxa, whereas most pathway resolved hits here were assimilatory or organic nitrogen genes, so direct comparison is limited. Relaxed threshold matches were retained for sensitivity assessment only and are not evidence of ecological or functional alteration (Zhao et al. 2024). Because most NCyc ORFs lacked pathway assignment and contig-based analysis cannot resolve host or activity, these results do not support inference that ARG carriage alters core benthic nitrogen-cycling processes. Analyzing these requires pathway resolved functional characterization and direct measurement of nitrogen-transformation activity (Kuypers et al. 2018).

### 3.6 Host taxonomic assignment and pathogen risk stratification

#### 3.6.1 Taxonomic distribution of ACCs

CAT assigned the 2,402 ACCs at variable taxonomic resolution, 217 reached species level (9.0%), 235 genus (9.8%), 83 family (3.5%), 116 order (4.8%), 242 class (10.1%) and 418 phylum (17.4%), while 1,091 (45.4%) had no supported rank (**Fig. S6; Table S18A**). Resolution differed among sites, with 70.9% of ACCs resolved to at least phylum at NRH and 63.3% at SM, compared with 34.2% at MM, where only one ACC reached species level. Species-level inference was therefore restricted to a small subset of the recovered resistome, and genus- or species-level host inference to 452 ACCs (18.8%) (**Table S18A**). This variable resolution is consistent with CAT’s use of multiple ORF-level homology signals to assign contigs at taxonomic ranks supported by available reference genomes, rather than forcing species-level classifications for poorly represented or uncharacterized sequences (Von Meijenfeldt et al. 2019). Consequently, host attribution for most ACCs remained limited to higher taxonomic ranks, constraining fine scale interpretation of resistome-host relationships (Guan et al. 2022).

Among the 1,309 ACCs classified to at least phylum rank, *Pseudomonadota* was the most frequent phylum (899 ACCs; 68.7%) (**Table S18E**). Recurrent ARG subtype-phylum associations (≥2 ACCs; 137 edges) were dominated by *Pseudomonadota*, including *mexB* (75 ACCs), *mexK* (73), *bacA* (70), *smeE* (63) and *novA* (50), each recovered across all six metagenomes. Additional associations included *muxB* (36), *mexW* (33), *rosB* (33), *mexF* (32) and *msbA* (28), each recovered at four to five sites (**Fig. 6A; Table S18B**). Secondary phylum-level contexts included *Actinomycetota*-associated *novA* on 27 ACCs from NH, NRH and SPH, and *Bacteroidota*-associated *β-lactamase* CGA-1 on 11 ACCs from both Adyar sites. The predominance of *Pseudomonadota* is consistent with previous metagenomic studies of river sediments that identified *Proteobacteria*/*Pseudomonadota* as major reservoirs of ARGs, particularly multidrug-resistance and efflux-associated determinants (Chen et al. 2019). At the family rank (66 recurrent edges), *Moraxellaceae* accounted for seven of the twelve highest-ranked edges, led by *mexB* (19 ACCs; MM and SM) and *macB* (11; MM, SM and NRH), followed by *tolC* (8), *abeM* (7), and *OprM*, *adeI* and *mexK* (6 each), CGA-1 localized to *Weeksellaceae* (11; MM and SM) (**Fig. S6; Table S18C**). *Acinetobacter* contained the largest set of recurrent genus-level associations (12 of 46 edges), with the seven most frequent, namely *mexB* (19), *macB* (10), *tolC* (8), *abeM* (7), and *OprM*, *adeI* and *mexK* (6 each), all supported at MM and SM (**Table S18D**).

The frequent recovery of efflux-associated genes (*mexB*, *mexK*, *OprM*, *macB*, *tolC*, *abeM*, and *adeI*) is consistent with the widespread occurrence of multidrug efflux machinery among Gram-negative bacterial lineages, where RND-type (*MexAB*-*OprM*-like) and *MacAB*-*TolC*-type systems constitute major antibiotic export mechanisms (Glavier et al. 2020; Lodé et al. 2025). Their distribution is also consistent with environmental observations that ARG profiles are strongly structured by microbial host composition (Chen et al. 2019). However, many multidrug transporter families are intrinsic chromosomal features of Gram-negative bacteria that mediate general physiological homeostasis, environmental stress adaptation and cellular detoxification in addition to antibiotic export. Sequence-level detection alone therefore cannot be interpreted as evidence of acquired, clinically transferable resistance without supporting genomic mobility, expression, or resistance phenotype (Nikaido 2009; Blanco et al. 2016). Heavily polluted sediments of the Mutha River in Pune, India showed enrichment of *Acinetobacter* populations and clinically relevant carbapenemase genes (*bla_OXA-58_*) under untreated urban wastewater influence (Marathe et al. 2017). *Acinetobacter*-associated loci recovered here were primarily efflux system annotations, including *adeI* and *abeM*, which are commonly regarded as intrinsic genomic features of *Acinetobacter*. This distinction cautions against transferring clinical interpretations between river systems, while neither proving chromosomal localization nor excluding clinically relevant resistance in the present samples (Nikaido 2009; Martínez et al. 2015).

#### 3.6.2 Human pathogenic bacteria context

Host taxonomy was placed in a public-health context using a descriptive, contig-level evidence scheme (C1-C5; **Table S19A**) that combines two criteria. i.e., correspondence of the CAT assignment with the human pathogenic bacteria (HPB) reference list of Zhou et al. (2019), at species level (confirmed) or genus level (suggestive), and genomic mobility evidence (geNomad plasmid-, virus- or provirus-like prediction, or an ARG-MGE pair within 5 kb. This scheme is conceptually related to ARG risk-ranking frameworks that integrate host pathogenicity and genomic mobility (Zhang et al. 2021), but it classifies contigs, not genes, and does not estimate infection probability or transmission risk from environmental metagenomic assemblies.

No ACC resolved to species level matched the HPB list, so no C1 or C2 classifications were obtained (**Table S19A**). Species-level assignment was available for only 217 of 2,402 ACCs (9.0%; **Table S18A and S19D**), and just three species-resolved ACCs fell within HPB-listed genera (*Pseudomonas mangrovi*, 2; *P. fluvialis*, 1), neither species being on the list (**Table S19E**). This should not be interpreted as evidence of pathogen absence, because many environmental lineages remain underrepresented in reference databases (Singh et al. 2026). Genus-level correspondence was detected on 127 ACCs (5.3% of all ACCs; 28.1% of the 452 ACCs resolved to genus or species), concentrated at the Adyar sites i.e., SM, 107 (13.6% of SM ACCs); MM, 19 (5.9%) and low at NRH, 1 (0.2%), with none at NH, MRH or SPH (**Fig. 8B; Table S19D**). These ACCs were assigned to *Acinetobacter* (83), *Pseudomonas* (27, all SM), *Aeromonas* (16, all SM) and *Corynebacterium* (1, MM) (**Fig. S8; Table S19B**), genera that include many non-pathogenic environmental species. Of the 127, 125 lacked mobility evidence (C4; 5.2% of all ACCs) (**Fig. 8B; Table S19A**). Two *Acinetobacter*-assigned ACCs combined genus-level correspondence with mobility evidence (C3a; 0.08% of ACCs; **Table S19C**). MM contig k141_98629 (3,398 bp) was plasmid-like (geNomad score 0.97) and carried a transposase (*tnpA*) ORF 2,810 bp from the ARG on a distinct ORF, whereas SM contig k141_111524 (4,353 bp) was plasmid-like (score 0.82) without any MGE ORF. A further 73 ACCs (3.0%) had mobility evidence without HPB-genus correspondence (C3), and 2,202 (91.7%) had neither (C5) (**Fig. S8; Table S19D**).

**Figure 8.**
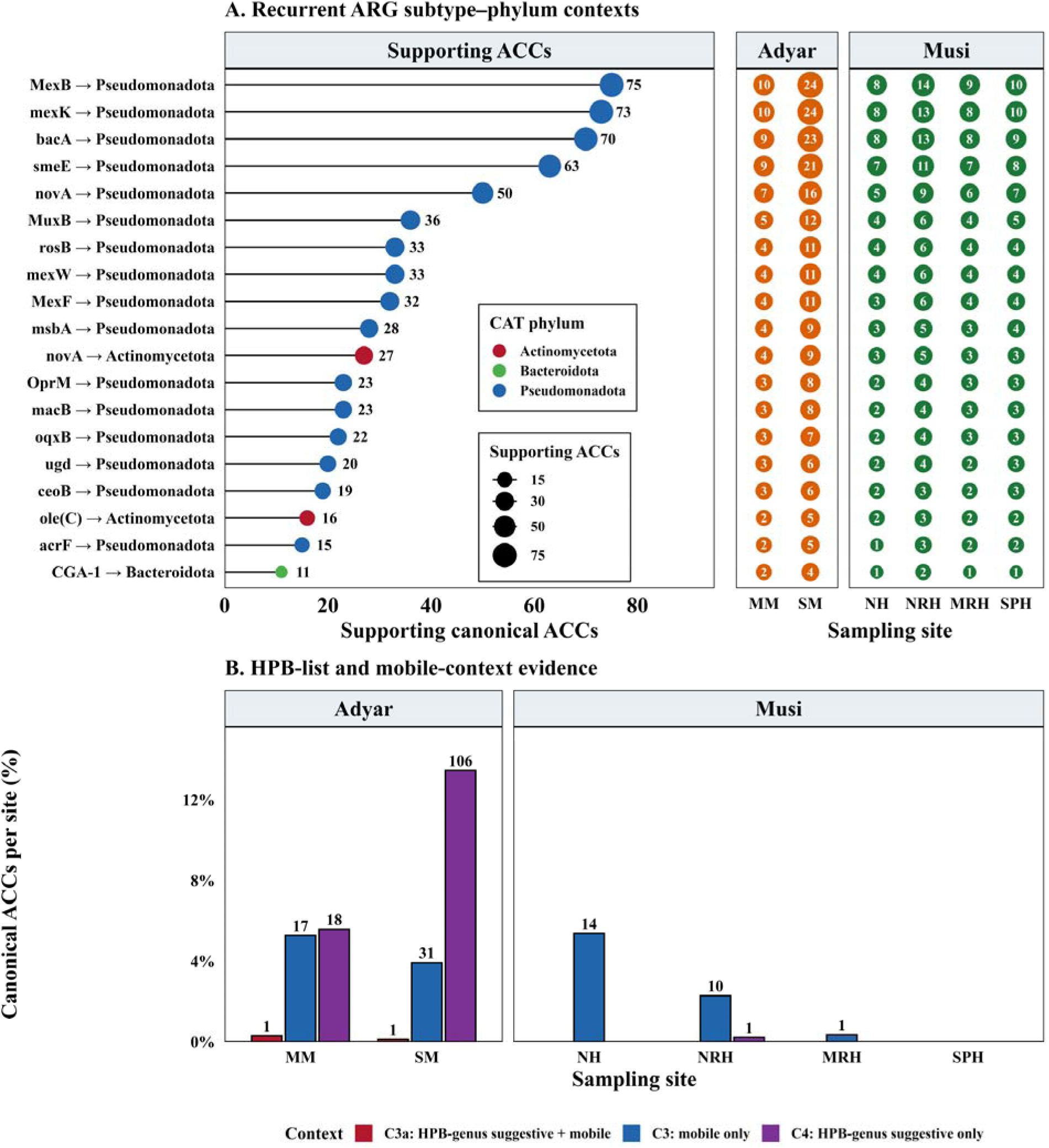
**(A)** Distribution of recurrent ARG subtype-phylum lineages, and **(B)** Mobile HPB risk classifications across Adyar and Musi River sites.

Species-level resolution was too limited to determine HPB carriage, but genus-level correspondence indicates that potentially pathogen-associated lineages account for a substantial fraction of the resolvable resistome at the Adyar sites (**Fig. S8; Table 19D**). Similar constraints affect other environmental metagenomic studies, where ARG-host relationships are often resolved only to higher ranks. In the Yangtze River, human pathogenic bacteria made up 13.4% and 5.9% of ARG hosts in water and sediment yet carried 64.4% and 46.1% of ARG subtypes (Wang et al. 2022). The present results are not a direct contradiction of that finding because the studies differ in sampling, taxonomic assignment and their ACCs were assigned to species by LCA against NCBI nr, and HPB also included virulence gene with different HPB definition and assembly characteristics. The C1-C5 scheme should therefore be viewed as a way of organizing host-correspondence and mobility evidence, not as an estimate of infection probability or sediment to human transmission (Zhang et al. 2021). Genus-level correspondence is only suggestive, since pathogenicity cannot be inferred from genus alone. The two C3a contigs are short, single-ARG contigs with moderate taxonomic support. They are potential candidates for host-resolved validation, such as long-read assembly or isolate-based confirmation, and not confirmed pathogen-dissemination hotspots. Mobility-associated ARGs need similar kind of validation before their epidemiological significance can be inferred (Tian et al. 2024). Together, these findings support a precautionary One Health interpretation while avoiding equivalence between mobile ARG detection and clinically relevant resistance.

## 5. Limitations

This study has few limitations that should be considered when interpreting the findings. Short-read metagenomic assembly can fragment mobile genetic contexts and constrain confident host attribution, potentially obscuring ARG-associated mobility (Arredondo-Alonso et al. 2017). Furthermore, ACC-based analyses reflect gene presence within ACCs rather than quantitative abundance or community-wide prevalence. Homology-based annotations against SARG, BacMet, VFDB, and NCyc databases indicate putative functional associations but do not establish gene expression, phenotypic resistance, host viability, or transmission potential (Karkman et al. 2018). Future investigations of urban river sediments should address these limitations through replicated spatial and seasonal sampling, integration of source tracking, sediment geochemistry, and the application of long-read or hybrid metagenomic sequencing. These complementary approaches could improve the resolution of linkages among ARGs, bacterial hosts, and mobile genetic elements. Additionally, isolate-based genomic characterization and antimicrobial susceptibility testing would provide independent evidence of viable ARG carriage and corresponding resistance phenotypes. Such methodological integration would strengthen risk-oriented environmental AMR assessments while avoiding the assumption that genomic detection alone establishes phenotypic resistance or public health risk (Mao et al. 2024; Li et al. 2026).

## 6. Conclusion

This study characterized ARG diversity, mobility-associated signatures, and putative bacterial hosts in wastewater-impacted sediments from two contrasting Indian river catchments. The resistomes exhibited substantial diversity and spatial heterogeneity, highlighting the importance of genomic context over ARG abundance or richness alone in assessing environmental resistance profiles. The co-occurrence of mobility-associated evidence and taxonomic correspondence with human pathogenic bacteria was limited. Instead, the benthic resistomes were predominantly characterized by efflux-associated and stress-response determinants, many of which are associated with intrinsic resistance mechanisms in Gram-negative bacteria. These findings underscore the importance of distinguishing environmental ARG detection from evidence of acquired or potentially transferable antimicrobial resistance. Although ARG detection alone does not establish a public health risk, the subset occurring in mobile or pathogen-associated genomic contexts warrants further investigation through long-read sequencing, host-resolved validation, and phenotypic characterization. Integrating these approaches with source tracking and sediment geochemistry could strengthen evidence-based surveillance of municipal sewage, hospital effluents, and other wastewater inputs while accounting for the uncertainty inherent in sequence-based risk assessment. Further evaluation across geographically and environmentally diverse catchments will be necessary to establish the broader applicability of this framework for comparative environmental AMR surveillance.

## CRediT authorship contribution statement

**Arun Kashyap:** Conceptualization, Methodology, Software, Formal analysis, Validation, Investigation, Data Curation, Writing - Original Draft, Writing - Review & Editing, Visualization. **Vikas Sonkar:** Methodology, Data Curation, Writing – review & editing, Investigation. **Indumathi M. Nambi**: Project Administration, Writing – review & editing. **Shashidhar Thatikonda:** Supervision, Formal analysis, Methodology, Project Administration, Conceptualization, Writing – review & editing, Validation, Visualization.

## Data availability

All data, including the additional figures and necessary tables are made available as part of supplementary information.

## Declaration of generative AI and AI-assisted technologies in the manuscript preparation process

During drafting of this study, the authors used Copilot to assist with English grammar corrections. After using them, the authors reviewed and edited the content as needed and take full responsibility for the content of the published article.

## Acknowledgements

We cheerfully acknowledge all laboratory associates for their invaluable assistance. We sincerely thank Dr. Sangeetha C. Jambu for reviewing this manuscript and assisting with manuscript refinement.

