## Supplementary Information for "Contig-Resolved Genetic Context of Antimicrobial Resistome in Sediments of Two Contrasting Wastewater-Impacted Rivers"

**Table S1.** Locations and coordinates of sediment sampling sites in the Adyar and Musi Rivers.

| **River** | **Site** | **SampleID** | **Latitude** | **Longitude** |
| --- | --- | --- | --- | --- |
| Adyar River, Tamil Nadu | Manimangalam | MM | 12.9324 | 80.0460 |
|  | Saidapet | SM | 13.0168 | 80.2248 |
| Musi River, Telangana | Nagole | NH | 17.3874 | 78.5724 |
|  | Nagaram Bridge | NRH | 17.3605 | 79.2717 |
|  | Musi Reservoir | MRH | 17.2323 | 79.5061 |
|  | Shunya Pahad | SPH | 16.7375 | 79.6680 |

**Table S2.** Sequence processing, gene catalog construction, mapping, and ARG reporting criteria

| **Analytical step** | **Software or method** | **Criterion** |
| --- | --- | --- |
| Minimum contig length | MEGAHIT v1.2.9 (Li et al. 2016) | ≥ 500 bp |
| ORF prediction | Prodigal v2.6.3 (Hyatt et al. 2010) | Metagenomic mode (-p meta) |
| Non-redundant protein clustering | CD-HIT v4.8.1 (Li and Godzik 2006) | 95% amino-acid identity; 90% coverage of the shorter sequence |
| CD-HIT identity calculation | CD-HIT | Local identity calculation (-G 0) |
| CD-HIT clustering mode | CD-HIT | Accurate greedy clustering (-g 1) |
| Read mapping | Bowtie2 v2.5.5 (Langmead and Salzberg 2012) | End-to-end sensitive mode |
| Paired-read insert interval | Bowtie2 | 0–1,000 bp |
| Mapping processing | SAMtools v1.23.1 (Danecek et al. 2021) | Mapped alignments retained, sorted, and indexed |
| Raw detection | Read mapping | ≥1 mapped read |
| Reportable ARG detection | Read mapping and breadth | ≥3 mapped reads and ≥50% ORF breadth |
| ARG abundance normalization | Coverage normalized by clean-data volume | Coverage ×/Gb |

**Table S3.** Reference databases, sequence-search methods, and filtering thresholds used to annotate ARGs and genomic context features.

| **Feature** | **Reference database** | **Query** | **Search method** | **Minimum identity** | **Minimum query coverage** | **E-value threshold** | **Additional filtering** | **Retained annotation** |
| --- | --- | --- | --- | --- | --- | --- | --- | --- |
| Antibiotic resistance genes (ARGs) | SARG v2.0 (Yin et al. 2018) | Predicted proteins | DIAMOND BLASTP (Buchfink et al. 2014), more-sensitive mode | 60% amino-acid identity | 70% | ≤1 × 10⁻¹⁰ | Maximum five initial target matches per ORF | One best-supported hit per ORF |
| Biocide and metal resistance genes (BMRGs) | BacMet2 (Pal et al. 2014) | Predicted proteins | DIAMOND BLASTP, more-sensitive mode | 70% amino-acid identity | 70% | ≤1 × 10⁻⁵ | Maximum five initial target matches per ORF | One best-supported hit per ORF |
| Virulence factor genes (VFGs) | VFDB (Zhou et al. 2025) | Predicted proteins | DIAMOND BLASTP, more-sensitive mode | 80% amino-acid identity | 70% | ≤1 × 10⁻⁵ | Maximum five initial target matches per ORF | One best-supported hit per ORF |
| Mobile genetic element-associated sequences (MGEs) | MGE (Pärnänen et al. 2018) | Representative nucleotide ORFs | BLASTN | 80% nucleotide identity | Not explicitly applied | ≤1 × 10⁻⁵ | Minimum alignment length of 75 bp; maximum five initial target matches per ORF | One best-supported hit per ORF |
| Nitrogen-cycling genes, primary screen | NCycDB 100% reference set (Tu et al. 2019) | Predicted proteins | DIAMOND BLASTP, more-sensitive mode | 85% amino-acid identity | 80% | ≤1 × 10⁻⁵ | Maximum five initial target matches per ORF | One best-supported hit per ORF |
| Nitrogen-cycling genes, relaxed screen | NCycDB 95% reference set (Tu et al. 2019) | Predicted proteins | DIAMOND BLASTP, more-sensitive mode | 70% amino-acid identity | 70% | ≤1 × 10⁻³ | Used as a sensitivity screen; maximum five initial target matches per ORF | One best-supported hit per ORF |

Notes: Query coverage refers to the proportion of the predicted ORF covered by the reference alignment. For each database, up to five candidate matches were initially returned. The best supported match per ORF was subsequently retained by ranking hits according to bit score, E-value, percentage identity, alignment length, and reference identifier. The MGE screen did not apply an explicit query-coverage threshold; instead, matches were required to have ≥80% nucleotide identity, an E-value ≤1 × 10⁻⁵, and an aligned length ≥75 bp. Relaxed NCyc annotations were retained for sensitivity assessment and were interpreted separately from primary annotations.

**Table S4.** Software, database, and parameters used for taxonomic annotation of ACC

| **Component** | **Specification** |
| --- | --- |
| Target sequences | Sample specific ARG-risk contigs |
| Taxonomic classifier | CAT pack, contigs workflow (Von Meijenfeldt et al. 2019) |
| Reference database | CAT-NR, release dated 12 December 2024 |
| Taxonomy | NCBI taxonomy files |
| Search sensitivity | Sensitive mode |
| Classification parameters | CAT pack defaults |
| Computational threads | 20 |
| Taxon-name processing | Official names added using CAT_pack add_names --only_official |

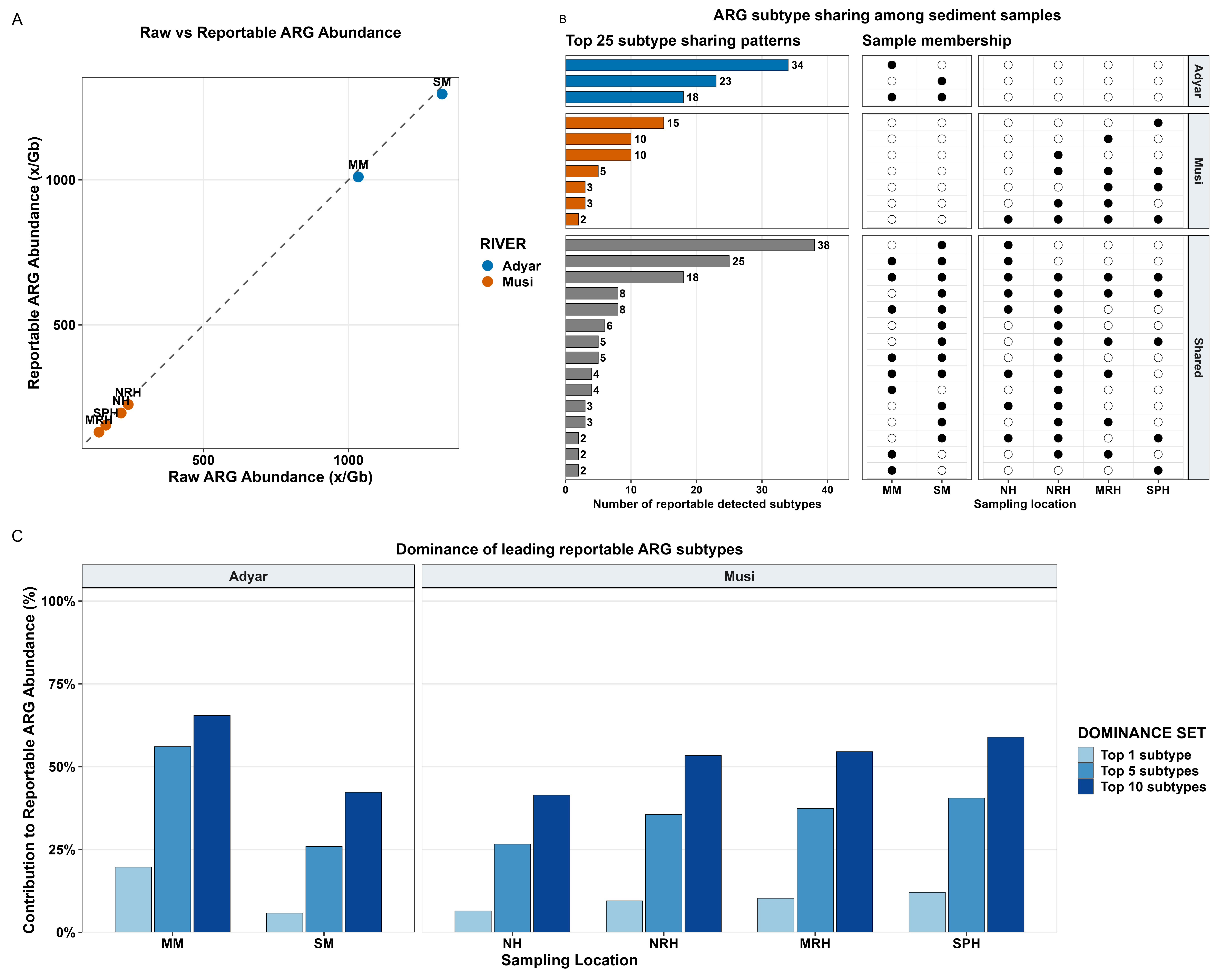

**Figure S1. (A)** Raw versus reportable ARG abundance, **(B)** subtype sharing patterns, and **(C)** subtype dominance across sediment metagenomes from the Adyar and Musi rivers.

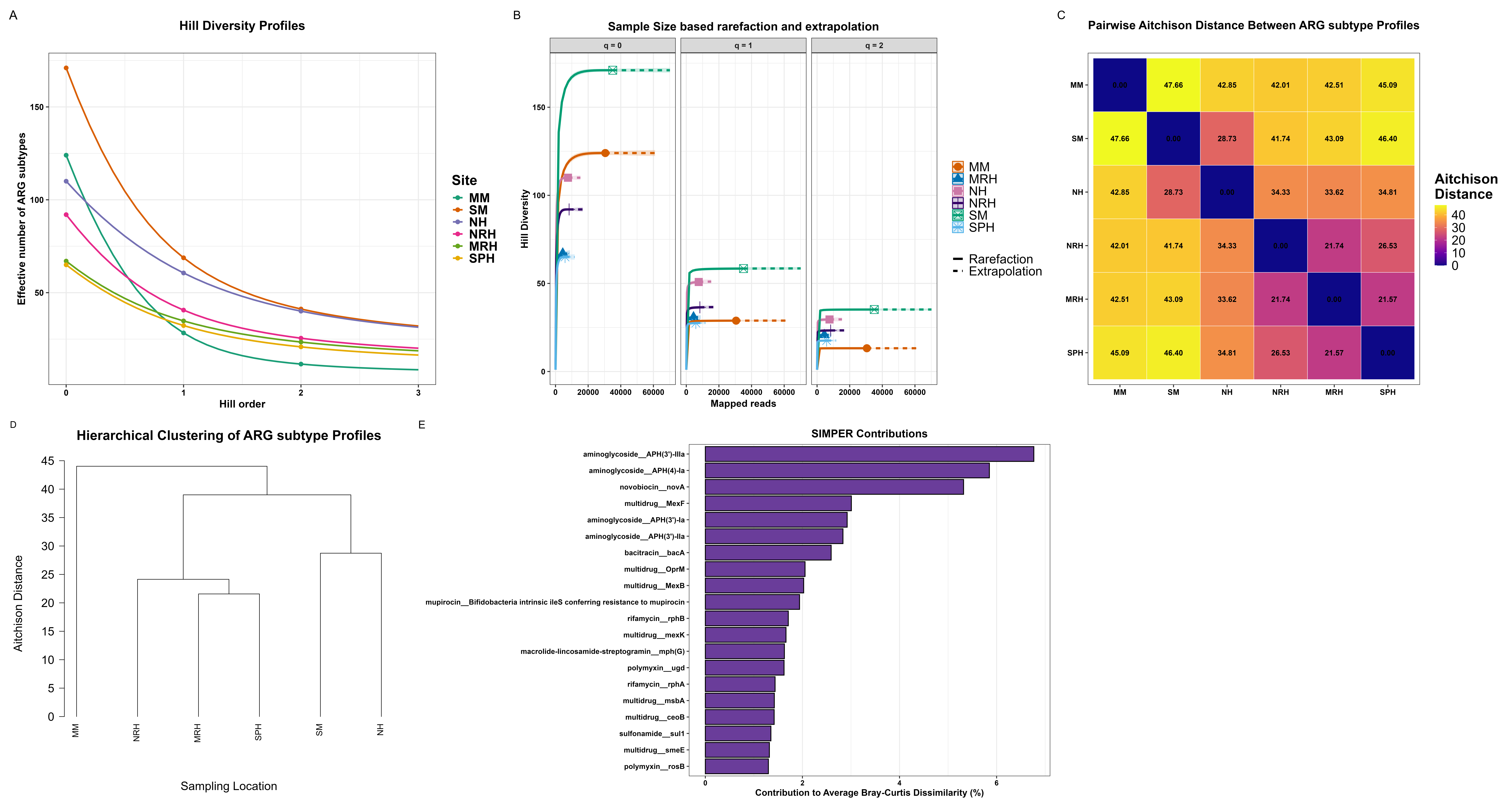

**Figure S2.** Hill diversity, sampling completeness and compositional analyses of ARG subtype profiles in the Adyar and Musi rivers. **(A)** Hill diversity profiles of ARG subtypes across sampling locations. **(B)** Sample-size-based rarefaction and extrapolation curves of ARG subtype richness. **(C)** Pairwise Aitchison distances between ARG subtype profiles. **(D)** Hierarchical clustering of ARG subtype profiles based on Aitchison distance. **(E)** ARG subtypes contributing most to river dissimilarity using SIMPER.

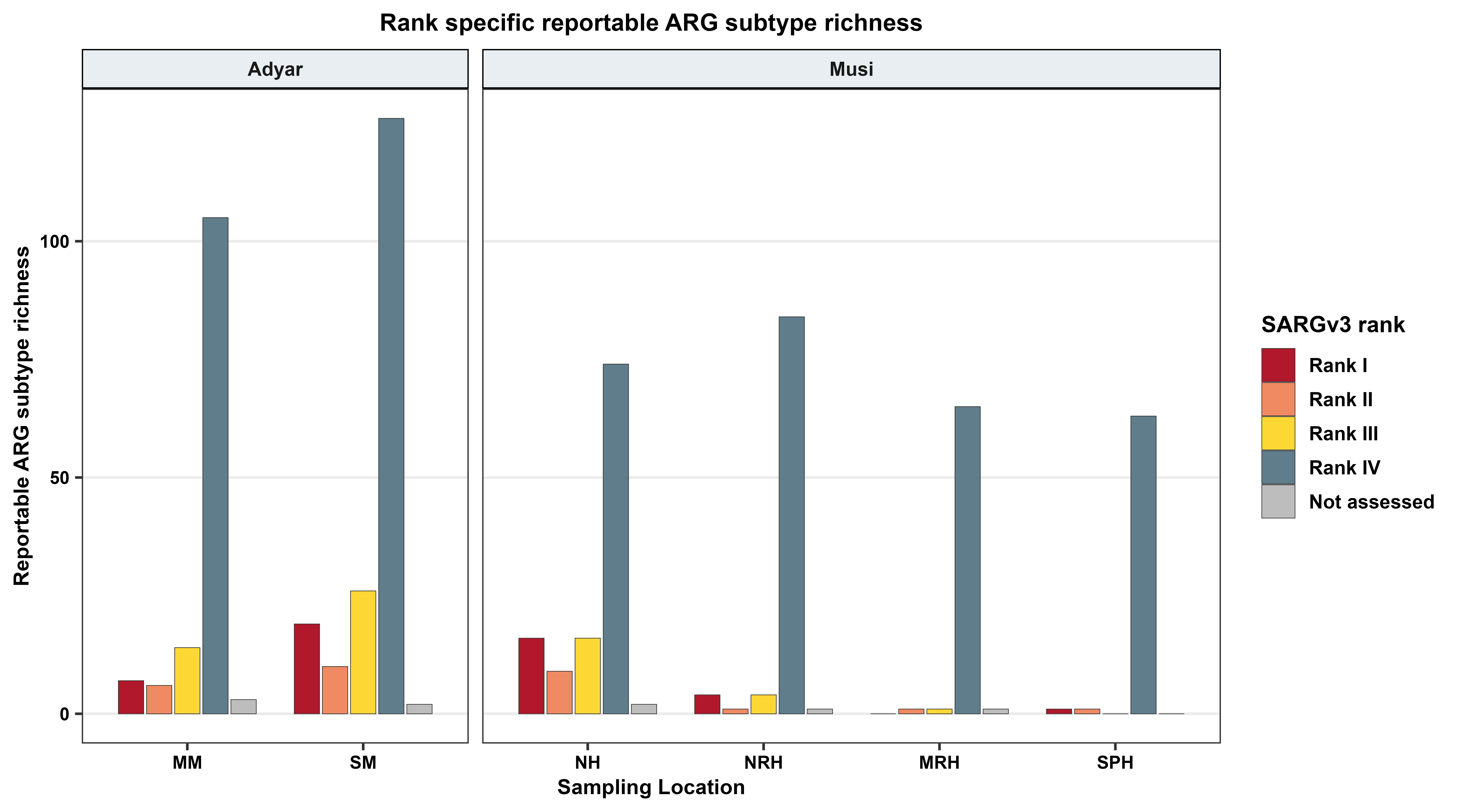
**Figure S3.** Rank-specific ARG subtype richness across sediment metagenomes. Number of reportable ARG subtypes in each SARGv3 rank (I–IV and not assessed;(Yin et al. 2023)) at each sampling location in the Adyar and Musi rivers.

**Table S5.** Reportable normalized ARG abundance (×/Gb) across sediment metagenomes categorized by SARGv3 risk rank .

| **SARGv3 rank** | **MM** | **SM** | **NH** | **NRH** | **MRH** | **SPH** |
| --- | --- | --- | --- | --- | --- | --- |
| Rank I | 212.630 | 64.529 | 19.536 | 1.062 | 0.000 | 0.258 |
| Rank II | 13.692 | 151.496 | 21.548 | 11.316 | 1.429 | 0.399 |
| Rank III | 23.027 | 93.127 | 22.358 | 1.945 | 0.339 | 0.000 |
| Rank IV | 754.820 | 969.586 | 128.546 | 209.480 | 128.550 | 153.898 |
| Not assessed | 5.923 | 17.157 | 4.401 | 1.871 | 0.155 | 0.000 |

**Table S6.** Summary and completeness metrics of SARGv3 risk-rank annotation for NR-ARG sequence representatives.

| **Metric** | **Value** |
| --- | --- |
| NR ARG representatives | 2055 |
| Unique SARG reference variants | 668 |
| Ranked NR-ARG representatives | 2055 |
| Unmatched NR-ARG representatives | 0 |
| Ranked SARG reference variants | 668 |
| Unmatched SARG reference variants | 0 |
| Static rank reference variants | 12746 |

**Table S7.** Reportable NR-ARG sequence representative counts (ORF-level richness) per sample categorized by SARGv3 risk rank.

| **SARGv3 rank** | **MM** | **SM** | **NH** | **NRH** | **MRH** | **SPH** |
| --- | --- | --- | --- | --- | --- | --- |
| Rank I | 7 | 27 | 22 | 4 | 0 | 1 |
| Rank II | 14 | 25 | 17 | 1 | 1 | 1 |
| Rank III | 18 | 48 | 30 | 4 | 1 | 0 |
| Rank IV | 316 | 588 | 248 | 385 | 275 | 290 |
| Not assessed | 3 | 2 | 2 | 1 | 1 | 0 |

**Table S8.** Distribution of mobility-associated genomic-context signals among ACC by sampling sites.

| **River** | **Location** | **ACCs** | **Plasmid-like** | **Virus-like** | **Provirus-regions** | **ARG-MGE** | **Any mobile** | **ARG overlaps provirus** | **Any mobile ACCs (%)** |
| --- | --- | --- | --- | --- | --- | --- | --- | --- | --- |
| Adyar | MM | 322 | 14 | 3 | 2 | 5 | 18 | 0 | 5.59 |
|  | SM | 787 | 20 | 3 | 0 | 17 | 32 | 0 | 4.07 |
| Musi | NH | 260 | 10 | 0 | 0 | 10 | 14 | 0 | 5.38 |
|  | NRH | 436 | 8 | 0 | 0 | 4 | 10 | 0 | 2.29 |
|  | MRH | 277 | 1 | 0 | 0 | 0 | 1 | 0 | 0.36 |
|  | SPH | 320 | 0 | 0 | 0 | 0 | 0 | 0 | 0 |

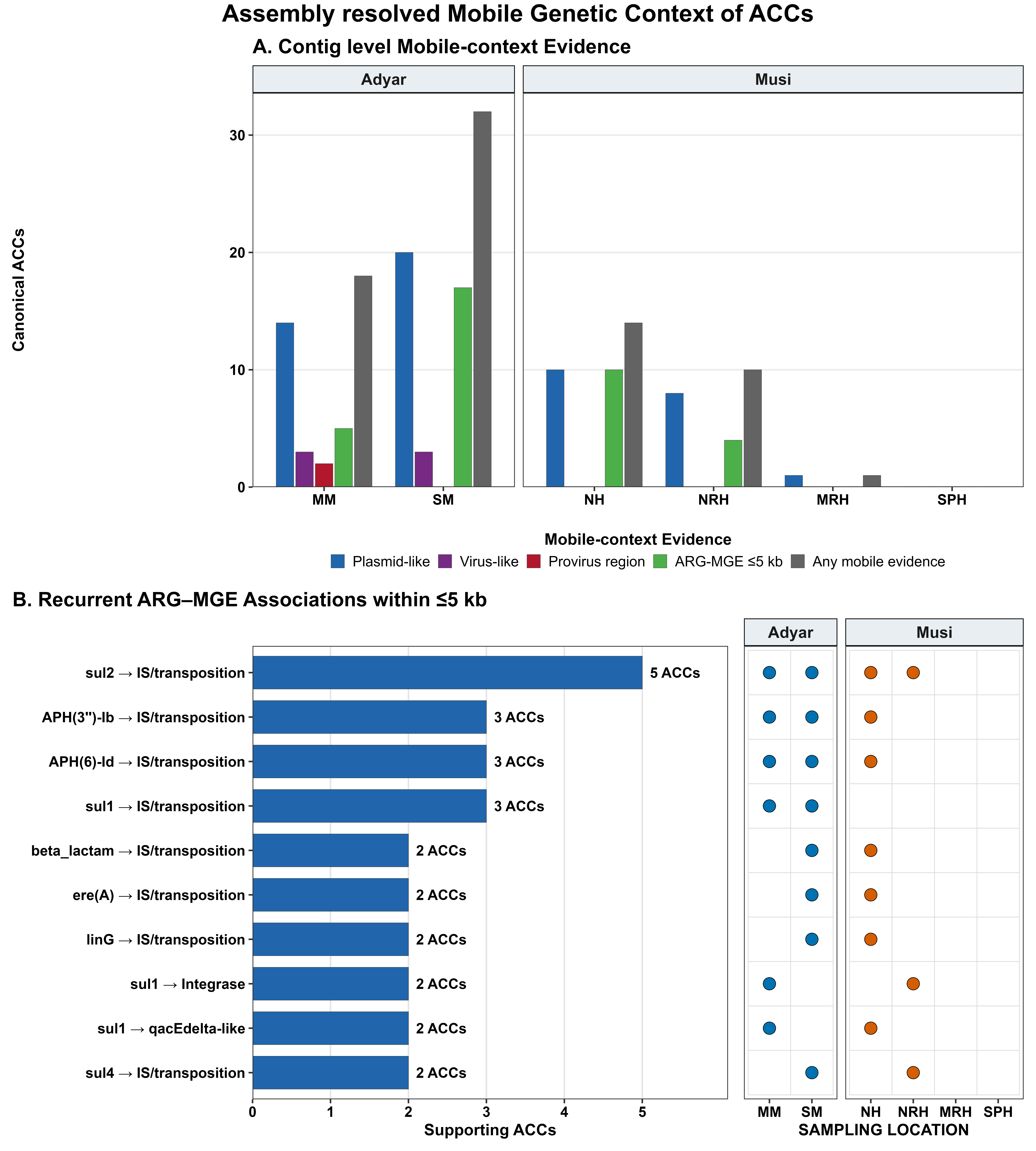
**Figure S4.** **(A)** Number of canonical ACCs per sample with plasmid-like, virus-like, provirus-region, ARG-MGE (≤ 5 kb) or any mobile evidence. **(B)** Recurrent ARG-MGE associations within ≤ 5 kb, supported by ≥ 2 canonical ACCs. Bars show the number of supporting ACCs, and filled circles mark the supporting samples across sediment metagenomes.

**Table S9.** Functional reclassification and site distribution of MGE reference annotations recovered on ACCs.

| **River** | **Location** | **Insertion sequence/transposition** | **Integrase** | ***qacEΔ*** |
| --- | --- | --- | --- | --- |
| Adyar | MM | 7 | 1 | 1 |
|  | SM | 16 | 1 | 5 |
| Musi | NH | 10 | 1 | 2 |
|  | NRH | 3 | 1 | 0 |
|  | MRH | 0 | 0 | 0 |
|  | SPH | 0 | 0 | 0 |

*qacEΔ*-like annotations were treated as integron-associated resistance markers rather than independent mobility machinery.

**Table S10.** Genomic distance summary for distinct ARG-like and MGE-like ORF pairs on ACCs.

| **ARG-MGE pairs** | **Pairs within 5kb** | **Pairs over 5kb** | **Median distance (bp)** | **Max distance (bp)** |
| --- | --- | --- | --- | --- |
| 54 | 52 | 2 | 806.5 | 94248 |

Pair counts exceed ACC counts because an ACC may contain more than one eligible ARG–MGE ORF combination.

**Table S11.** Recurrent ARG subtype-MGE category associations supported by at least two ACCs.

| **ARG subtype** | **Target** | **ACCs** | **Locations** |
| --- | --- | --- | --- |
| sulfonamide (*sul2*) | IS/Tp | 5 | MM;NH;NRH;SM |
| aminoglycoside (*APH(3’’)-Ib*) | IS/Tp | 3 | MM;NH;SM |
| aminoglycoside (*APH(6)-Id*) | IS/Tp | 3 | MM;NH;SM |
| sulfonamide (*sul1*) | IS/Tp | 3 | MM;SM |
| beta lactam (*class A beta-lactamase*) | IS/Tp | 2 | NH;SM |
| MLS *(ere(A))* | IS/Tp | 2 | NH;SM |
| MLS (*linG*) | IS/Tp | 2 | NH;SM |
| sulfonamide (*sul1*) | integrase | 2 | MM;NRH |
| sulfonamide (*sul1*) | *qacEΔ* | 2 | MM;NH |
| sulfonamide (*sul4*) | IS/Tp | 2 | NRH;SM |

MLS = Macrolide-Lincosamide-Streptogramin; IS/Tp = Insertion sequence or transposition.

**Table S12.** Site-level ACC support for recurrent ARG-MGE associations within 5 kb.

| **ARG subtype** | **MGE broad category** | **sample** | **ACCs site** | **River** | **ACCs** | **samples** |
| --- | --- | --- | --- | --- | --- | --- |
| aminoglycoside (*APH(3’’)-Ib*) | IS/Tp | SM | 1 | Adyar | 3 | MM;NH;SM |
| aminoglycoside (*APH(3’’)-Ib*) | IS/Tp | MM | 1 | Adyar | 3 | MM;NH;SM |
| aminoglycoside (*APH(3’’)-Ib*) | IS/Tp | NH | 1 | Musi | 3 | MM;NH;SM |
| aminoglycoside (*APH(3’’)-Ib*) | IS/Tp | NRH | 0 | Musi | 3 | MM;NH;SM |
| aminoglycoside (*APH(3’’)-Ib*) | IS/Tp | MRH | 0 | Musi | 3 | MM;NH;SM |
| aminoglycoside (*APH(3’’)-Ib*) | IS/Tp | SPH | 0 | Musi | 3 | MM;NH;SM |
| aminoglycoside (*APH(6)-Id*) | IS/Tp | SM | 1 | Adyar | 3 | MM;NH;SM |
| aminoglycoside (*APH(6)-Id*) | IS/Tp | MM | 1 | Adyar | 3 | MM;NH;SM |
| aminoglycoside (*APH(6)-Id*) | IS/Tp | NH | 1 | Musi | 3 | MM;NH;SM |
| aminoglycoside (*APH(6)-Id*) | IS/Tp | NRH | 0 | Musi | 3 | MM;NH;SM |
| aminoglycoside (*APH(6)-Id*) | IS/Tp | MRH | 0 | Musi | 3 | MM;NH;SM |
| aminoglycoside (*APH(6)-Id*) | IS/Tp | SPH | 0 | Musi | 3 | MM;NH;SM |
| beta lactam (*Other class A beta-lactamase*) | IS/Tp | SM | 1 | Adyar | 2 | NH;SM |
| beta lactam (*Other class A beta-lactamase*) | IS/Tp | MM | 0 | Adyar | 2 | NH;SM |
| beta lactam (*Other class A beta-lactamase*) | IS/Tp | NH | 1 | Musi | 2 | NH;SM |
| beta lactam (*Other class A beta-lactamase*) | IS/Tp | NRH | 0 | Musi | 2 | NH;SM |
| beta lactam (*Other class A beta-lactamase*) | IS/Tp | MRH | 0 | Musi | 2 | NH;SM |
| beta lactam (*Other class A beta-lactamase*) | IS/Tp | SPH | 0 | Musi | 2 | NH;SM |
| MLS *(ere(A))* | IS/Tp | SM | 1 | Adyar | 2 | NH;SM |
| MLS *(ere(A))* | IS/Tp | MM | 0 | Adyar | 2 | NH;SM |
| MLS *(ere(A))* | IS/Tp | NH | 1 | Musi | 2 | NH;SM |
| MLS *(ere(A))* | IS/Tp | NRH | 0 | Musi | 2 | NH;SM |
| MLS *(ere(A))* | IS/Tp | MRH | 0 | Musi | 2 | NH;SM |
| MLS *(ere(A))* | IS/Tp | SPH | 0 | Musi | 2 | NH;SM |
| MLS (*linG*) | IS/Tp | SM | 1 | Adyar | 2 | NH;SM |
| MLS (*linG*) | IS/Tp | MM | 0 | Adyar | 2 | NH;SM |
| MLS (*linG*) | IS/Tp | NH | 1 | Musi | 2 | NH;SM |
| MLS (*linG*) | IS/Tp | NRH | 0 | Musi | 2 | NH;SM |
| MLS (*linG*) | IS/Tp | MRH | 0 | Musi | 2 | NH;SM |
| MLS (*linG*) | IS/Tp | SPH | 0 | Musi | 2 | NH;SM |
| sulfonamide (*sul1*) | IS/Tp | SM | 2 | Adyar | 3 | MM;SM |
| sulfonamide (*sul1*) | IS/Tp | MM | 1 | Adyar | 3 | MM;SM |
| sulfonamide (*sul1*) | IS/Tp | NH | 0 | Musi | 3 | MM;SM |
| sulfonamide (*sul1*) | IS/Tp | NRH | 0 | Musi | 3 | MM;SM |
| sulfonamide (*sul1*) | IS/Tp | MRH | 0 | Musi | 3 | MM;SM |
| sulfonamide (*sul1*) | IS/Tp | SPH | 0 | Musi | 3 | MM;SM |
| sulfonamide (*sul1*) | integrase | SM | 0 | Adyar | 2 | MM;NRH |
| sulfonamide (*sul1*) | integrase | MM | 1 | Adyar | 2 | MM;NRH |
| sulfonamide (*sul1*) | integrase | NH | 0 | Musi | 2 | MM;NRH |
| sulfonamide (*sul1*) | integrase | NRH | 1 | Musi | 2 | MM;NRH |
| sulfonamide (*sul1*) | integrase | MRH | 0 | Musi | 2 | MM;NRH |
| sulfonamide (*sul1*) | integrase | SPH | 0 | Musi | 2 | MM;NRH |
| sulfonamide (*sul1*) | *qacEΔ* | SM | 0 | Adyar | 2 | MM;NH |
| sulfonamide (*sul1*) | *qacEΔ* | MM | 1 | Adyar | 2 | MM;NH |
| sulfonamide (*sul1*) | *qacEΔ* | NH | 1 | Musi | 2 | MM;NH |
| sulfonamide (*sul1*) | *qacEΔ* | NRH | 0 | Musi | 2 | MM;NH |
| sulfonamide (*sul1*) | *qacEΔ* | MRH | 0 | Musi | 2 | MM;NH |
| sulfonamide (*sul1*) | *qacEΔ* | SPH | 0 | Musi | 2 | MM;NH |
| sulfonamide *(sul2)* | IS/Tp | SM | 1 | Adyar | 5 | MM;NH;NRH;SM |
| sulfonamide *(sul2)* | IS/Tp | MM | 1 | Adyar | 5 | MM;NH;NRH;SM |
| sulfonamide *(sul2)* | IS/Tp | NH | 1 | Musi | 5 | MM;NH;NRH;SM |
| sulfonamide *(sul2)* | IS/Tp | NRH | 2 | Musi | 5 | MM;NH;NRH;SM |
| sulfonamide *(sul2)* | IS/Tp | MRH | 0 | Musi | 5 | MM;NH;NRH;SM |
| sulfonamide *(sul2)* | IS/Tp | SPH | 0 | Musi | 5 | MM;NH;NRH;SM |
| sulfonamide (*sul4*) | IS/Tp | SM | 1 | Adyar | 2 | NRH;SM |
| sulfonamide (*sul4*) | IS/Tp | MM | 0 | Adyar | 2 | NRH;SM |
| sulfonamide (*sul4*) | IS/Tp | NH | 0 | Musi | 2 | NRH;SM |
| sulfonamide (*sul4*) | IS/Tp | NRH | 1 | Musi | 2 | NRH;SM |
| sulfonamide (*sul4*) | IS/Tp | MRH | 0 | Musi | 2 | NRH;SM |
| sulfonamide (*sul4*) | IS/Tp | SPH | 0 | Musi | 2 | NRH;SM |

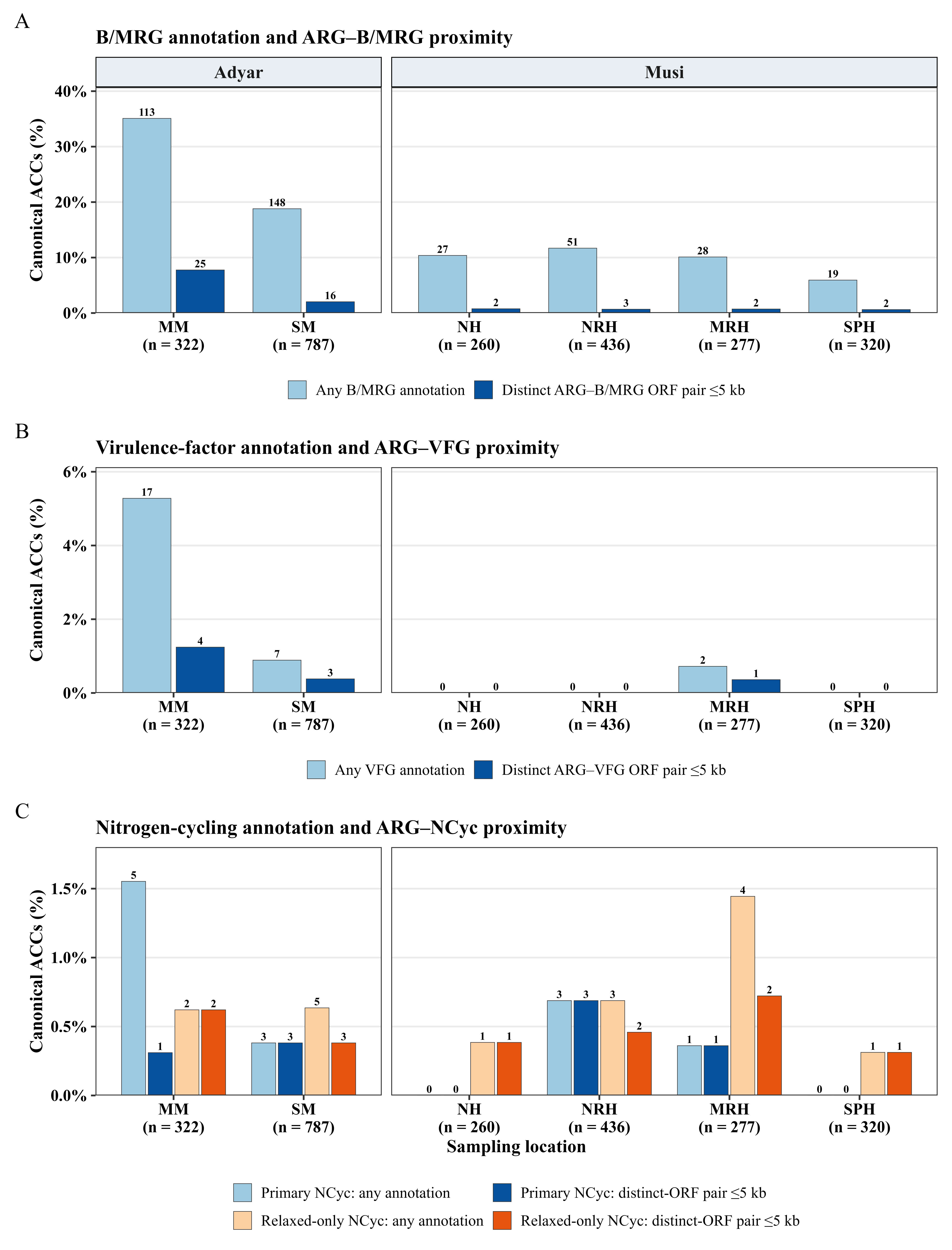
**Figure S5.** Site-level distribution of **(A)** BMRG, **(B)** VFG, and **(C)** nitrogen-cycling contexts among ACCs of Adyar and Musi metagenomes.

**Table S13.** Site-level occurrence and genomic relationships of BMRG annotations on ACCs.

| **River** | **Site** | **Total ACCs** | **ACCs with any B/MRG annotation** | **% of site ACCs (any B/MRG)** | **ACCs with same-ORF ARG-B/MRG annotation** | **% of site ACCs (same-ORF)** | **ACCs with distinct-ORF ARG-B/MRG (≤ 5 kb)** | **% of site ACCs (distinct-ORF ≤ 5 kb)** | **% of B/MRG-positive ACCs with distinct-ORF (≤ 5 kb)** |
| --- | --- | --- | --- | --- | --- | --- | --- | --- | --- |
| Adyar | MM | 322 | 113 | 35.1 | 105 | 32.6 | 31 | 9.6 | 27.4 |
|  | SM | 787 | 148 | 18.8 | 147 | 18.7 | 20 | 2.5 | 13.5 |
| Musi | NH | 260 | 27 | 10.4 | 27 | 10.4 | 3 | 1.2 | 11.1 |
|  | NRH | 436 | 51 | 11.7 | 51 | 11.7 | 3 | 0.7 | 5.9 |
|  | MRH | 277 | 28 | 10.1 | 28 | 10.1 | 2 | 0.7 | 7.1 |
|  | SPH | 320 | 19 | 5.9 | 19 | 5.9 | 2 | 0.6 | 10.5 |
| Total | All sites | 2402 | 386 | 16.1 | 377 | 15.7 | 61 | 2.5 | 15.8 |

**Table S14.** ARG subtype and BMRG gene associations recovered on the ACC within 5 kb.

| **ARG subtype** | **BMRG** | **ACC** | **Location** |
| --- | --- | --- | --- |
| multidrug *adeI* | *adeJ* | 6 | MM;NRH;SM |
| multidrug *MuxC* | *mdtB/yegN* | 4 | MM;NRH;SPH |
| multidrug *smeE* | *adeI* | 4 | MM;NRH;SM |
| multidrug *smeD* | *smeE* | 3 | MM;SM |
| multidrug *MexC* | *mexD* | 2 | NH;SM |
| multidrug *MexD* | *cnrR/cnrX* | 2 | NH;SM |
| multidrug *MuxB* | *mdtC/yegO* | 2 | MM;SPH |
| multidrug *OprJ* | *mexD* | 2 | NH;SM |
| sulfonamide *sul1* | *qacEdelta1* | 2 | MM;NH |
| multidrug *MexB* | *adeK* | 2 | SM |
| multidrug *MuxB* | *mdtC* | 2 | MM |
| multidrug *emrA* | *emrB* | 2 | MM |
| multidrug *emrA* | *emrR* | 2 | MM |
| multidrug *emrB* | *emrA* | 2 | MM |
| multidrug *mdtE* | *baeS* | 2 | MM |
| multidrug *mexV* | *mexW* | 2 | SM |
| multidrug *oqxA* | *oqxB* | 2 | MM |
| multidrug *oqxB* | *oqxA* | 2 | MM |
| multidrug *smeE* | *adeK* | 2 | MM |
| aminoglycoside *AAC(6')-Ib4* | *qacF* | 1 | SM |
| aminoglycoside *APH(3'')-Ib* | *merR* | 1 | MM |
| aminoglycoside *APH(6)-Id* | *merR* | 1 | MM |
| beta lactam *CMH-1* | *sugE* | 1 | MM |
| beta lactam *OXA-4* | *qacEdelta1* | 1 | SM |
| multidrug *Enterobacter cloacae acrA* | *acrB* | 1 | MM |
| multidrug *Enterobacter cloacae acrA* | *acrR/ybaH* | 1 | MM |
| multidrug *Escherichia coli acrA* | *sdeY* | 1 | MM |
| multidrug *MexA* | *emhR* | 1 | SM |
| multidrug *MexA* | *sdeY* | 1 | SM |
| multidrug *MexB* | *adeI* | 1 | SM |
| multidrug *MexB* | *eefA* | 1 | MM |
| multidrug *MexC* | *cnrR/cnrX* | 1 | NH |
| multidrug *MexE* | *adeG* | 1 | SPH |
| multidrug *MexE* | *mdtF/yhiV* | 1 | NH |
| multidrug *OprM* | *bepE* | 1 | SM |
| multidrug *abeM* | *abeM* | 1 | MM |
| multidrug *acrE* | *acrF/envD* | 1 | MM |
| multidrug *acrF* | *acrA* | 1 | MM |
| multidrug *acrF* | *acrE/envC* | 1 | MM |
| multidrug *acrF* | *fabI* | 1 | MM |
| multidrug *adeI* | *adeK* | 1 | MM |
| multidrug *ceoB* | *adeG* | 1 | MRH |
| multidrug *emrD* | *glpF* | 1 | MM |
| multidrug *mdtE* | *baeR* | 1 | MM |
| multidrug *mdtE* | *mdtC* | 1 | MM |
| multidrug *mdtE* | *mdtC/yegO* | 1 | MM |
| multidrug *mdtF* | *adeI* | 1 | SM |
| multidrug *mdtF* | *oscA* | 1 | SM |
| multidrug *mdtK* | *emmdR* | 1 | MM |
| multidrug *mexJ* | *mexK* | 1 | SM |
| multidrug *mexK* | *pcm* | 1 | MM |
| multidrug *mexK* | *tolCsm* | 1 | MM |
| multidrug *msbA* | *smrA* | 1 | NRH |
| multidrug *oqxA* | *sdeB* | 1 | MM |
| multidrug *sdeY* | *sdeX* | 1 | MM |
| multidrug *smeD* | *mdtF/yhiV* | 1 | SM |
| multidrug *smeD* | *sdeY* | 1 | MRH |
| multidrug *smeD* | *smeT* | 1 | MM |
| multidrug *smeE* | *oscA* | 1 | MM |
| multidrug *smeF* | *smeE* | 1 | SM |

**Table S15A.** Genomic co-localization of ARG subtypes biocide classes within 5 kb on the same contig that were supported by at least two ACCs.

| **ARG subtype** | **Biocide** | **ACCs** | **Location** |
| --- | --- | --- | --- |
| multidrug *adeI* | Acridine | 6 | MM;NRH;SM |
| multidrug *ade*I | Azin | 6 | MM;NRH;SM |
| multidrug *ade*I | Organo-sulfate | 6 | MM;NRH;SM |
| multidrug *adeI* | Xanthene | 6 | MM;NRH;SM |
| multidrug *smeD* | Organo-sulfate | 5 | MM;MRH;SM |
| multidrug *smeD* | Phenolic compounds | 5 | MM;MRH;SM |
| multidrug *smeD* | Xanthene | 5 | MM;MRH;SM |
| multidrug *smeE* | Acridine | 5 | MM;NRH;SM |
| multidrug *smeE* | Azin | 5 | MM;NRH;SM |
| multidrug *smeE* | Organo-sulfate | 5 | MM;NRH;SM |
| multidrug *smeE* | Xanthene | 5 | MM;NRH;SM |
| multidrug *MuxC* | Acid | 4 | MM;NRH;SPH |
| multidrug *smeD* | Acridine | 4 | MM;MRH;SM |
| multidrug *smeE* | Acid | 4 | MM;NRH;SM |
| multidrug *smeE* | Biguanides | 4 | MM;NRH;SM |
| multidrug *smeE* | Diamidine | 4 | MM;NRH;SM |
| multidrug *smeE* | Paraquat | 4 | MM;NRH;SM |
| multidrug *smeE* | Peroxysulfate | 4 | MM;NRH;SM |
| multidrug *smeE* | Phenanthridine | 4 | MM;NRH;SM |
| multidrug *smeE* | Phenyl | 4 | MM;NRH;SM |
| multidrug *smeE* | QAC-Glutaraldehyde | 4 | MM;NRH;SM |
| multidrug *smeE* | Quaternary Ammonium Compounds (QACs) | 4 | MM;NRH;SM |
| multidrug *smeE* | Triarylmethane | 4 | MM;NRH;SM |
| multidrug *smeD* | Triarylmethane | 4 | MM;SM |
| multidrug *emrA* | Hydrazone | 4 | MM |
| multidrug smeD | Phenanthridine | 3 | MM;SM |
| multidrug *MexB* | Acridine | 3 | SM |
| multidrug *MexB* | Azin | 3 | SM |
| multidrug *MexB* | Organo-sulfate | 3 | SM |
| multidrug *MexB* | Xanthene | 3 | SM |
| multidrug mdtE | Acid | 3 | MM |
| multidrug *oqxA* | Biguanides | 3 | MM |
| multidrug *oqxA* | Organo-sulfate | 3 | MM |
| multidrug *oqxA* | Quaternary Ammonium Compounds (QACs) | 3 | MM |
| multidrug *MexB* | Acid | 2 | MM;SM |
| multidrug *MexC* | Alkane | 2 | NH;SM |
| multidrug *MexC* | Aromatic hydrocarbons | 2 | NH;SM |
| multidrug *MexC* | Phenolic compounds | 2 | NH;SM |
| multidrug *MexE* | Organo-sulfate | 2 | NH;SPH |
| multidrug *MuxB* | Acid | 2 | MM;SPH |
| multidrug *OprJ* | Alkane | 2 | NH;SM |
| multidrug *OprJ* | Aromatic hydrocarbons | 2 | NH;SM |
| multidrug *OprJ* | Phenolic compounds | 2 | NH;SM |
| multidrug *smeD* | Quaternary Ammonium Compounds (QACs) | 2 | MRH;SM |
| sulfonamide *sul1* | Acid | 2 | MM;NH |
| sulfonamide *sul1* | Acridine | 2 | MM;NH |
| sulfonamide *sul1* | Biguanides | 2 | MM;NH |
| sulfonamide *sul1* | Diamindine | 2 | MM;NH |
| sulfonamide *sul1* | Organo-sulfate | 2 | MM;NH |
| sulfonamide *sul1* | Paraquat | 2 | MM;NH |
| sulfonamide *sul1* | Phenanthridine | 2 | MM;NH |
| sulfonamide *sul1* | Quaternary Ammonium Compounds (QACs) | 2 | MM;NH |
| sulfonamide *sul1* | Triarylmethane | 2 | MM;NH |
| sulfonamide *sul1* | Xanthene | 2 | MM;NH |
| multidrug *Enterobacter cloacae acrA* | Acridine | 2 | MM |
| multidrug *MexA* | Organo-sulfate | 2 | SM |
| multidrug *acrF* | Acridine | 2 | MM |
| multidrug *acrF* | Phenolic compounds | 2 | MM |
| multidrug *emrA* | Hydrazine | 2 | MM |
| multidrug *emrA* | Organo-mercury | 2 | MM |
| multidrug *emrB* | Hydrazine | 2 | MM |
| multidrug *emrB* | Hydrazone | 2 | MM |
| multidrug *emrB* | Organo-mercury | 2 | MM |
| multidrug *emrB* | Salicylanilide | 2 | MM |
| multidrug *mexV* | Acridine | 2 | SM |
| multidrug *mexV* | Phenanthridine | 2 | SM |
| multidrug *oqxA* | Acridine | 2 | MM |
| multidrug *oqxA* | Halogens | 2 | MM |
| multidrug *oqxA* | Phenolic compounds | 2 | MM |
| multidrug *oqxB* | Acridine | 2 | MM |
| multidrug *oqxB* | Biguanides | 2 | MM |
| multidrug *oqxB* | Halogens | 2 | MM |
| multidrug *oqxB* | Organo-sulfate | 2 | MM |
| multidrug *oqxB* | Phenolic compounds | 2 | MM |
| multidrug *oqxB* | Quaternary Ammonium Compounds (QACs) | 2 | MM |

**Table S15B.** Genomic co-localization of ARG subtypes metal classes within 5 kb on the same contig that were supported by at least two ACCs.

| **ARG subtype** | **Metal** | **ACCs** | **Location** |
| --- | --- | --- | --- |
| multidrug *mdtE* | Zinc | 3 | MM |
| multidrug *MexD* | Cobalt | 2 | NH;SM |
| multidrug *MexD* | Nickel | 2 | NH;SM |
| multidrug *MuxB* | Zinc | 2 | MM |
| multidrug *emrA* | Mercury | 2 | MM |
| multidrug *emrB* | Mercury | 2 | MM |

**Table S15C.** Genomic co-localization of ARG subtypes BMRG within 5 kb on the same contig that were supported by at least two ACCs.

| **ARG subtypes** | **BMRGs** | **ACCs** | **Location** |
| --- | --- | --- | --- |
| multidrug *adeI* | *adeJ* | 6 | MM;NRH;SM |
| multidrug *MuxC* | *mdtB/yegN* | 4 | MM;NRH;SPH |
| multidrug *smeE* | *adeI* | 4 | MM;NRH;SM |
| multidrug *smeD* | *smeE* | 3 | MM;SM |
| multidrug *MexC* | *mexD* | 2 | NH;SM |
| multidrug *MexD* | *cnrR/cnrX* | 2 | NH;SM |
| multidrug *MuxB* | *mdtC/yegO* | 2 | MM;SPH |
| multidrug *OprJ* | *mexD* | 2 | NH;SM |
| sulfonamide *sul1* | *qacEdelta1* | 2 | MM;NH |
| multidrug *MexB* | *adeK* | 2 | SM |
| multidrug *MuxB* | *mdtC* | 2 | MM |
| multidrug *emrA* | *emrB* | 2 | MM |
| multidrug *emrA* | *emrR* | 2 | MM |
| multidrug *emrB* | *emrA* | 2 | MM |
| multidrug *mdtE* | *baeS* | 2 | MM |
| multidrug *mexV* | *mexW* | 2 | SM |
| multidrug *oqxA* | *oqxB* | 2 | MM |
| multidrug *oqxB* | *oqxA* | 2 | MM |
| multidrug *smeE* | *adeK* | 2 | MM |

**Table S16A.** Prevalence of ACCs with VFDB matches per site.

| **River** | **Location** | **Total ACCs** | **ACCs with VF** | **ACCs with VF (%)** | **VF ORFs** | **same ORF as ARG** | **distinct ORF** |
| --- | --- | --- | --- | --- | --- | --- | --- |
| Adyar | MM | 322 | 17 | 5.28 | 19 | 19 | 0 |
|  | SM | 787 | 7 | 0.89 | 7 | 6 | 1 |
| Musi | NH | 260 | 0 | 0 | 0 | 0 | 0 |
|  | NRH | 436 | 0 | 0 | 0 | 0 | 0 |
|  | MRH | 277 | 2 | 0.72 | 2 | 2 | 0 |
|  | SPH | 320 | 0 | 0 | 0 | 0 | 0 |

**Table S16B.** ORF-level VFDB annotations (one accepted hit per ORF) with the closest ARG relationship.

| **Location** | **VF gene** | **VF category** | **Relationship** | **Distance (bp)** |
| --- | --- | --- | --- | --- |
| MM | *acrA* | Antimicrobial activity/Competitive advantage | Same ORF as ARG | 0 |
|  | *acrA* | Antimicrobial activity/Competitive advantage | Same ORF as ARG | 0 |
|  | *acrA* | Antimicrobial activity/Competitive advantage | Same ORF as ARG | 0 |
|  | *acrA* | Antimicrobial activity/Competitive advantage | Same ORF as ARG | 0 |
|  | *acrB* | Antimicrobial activity/Competitive advantage | Same ORF as ARG | 0 |
|  | *acrB* | Antimicrobial activity/Competitive advantage | Same ORF as ARG | 0 |
|  | *acrB* | Antimicrobial activity/Competitive advantage | Same ORF as ARG | 0 |
|  | *acrB* | Antimicrobial activity/Competitive advantage | Same ORF as ARG | 0 |
|  | *acrB* | Antimicrobial activity/Competitive advantage | Same ORF as ARG | 0 |
|  | *acrB* | Antimicrobial activity/Competitive advantage | Same ORF as ARG | 0 |
|  | *acrB* | Antimicrobial activity/Competitive advantage | Same ORF as ARG | 0 |
|  | *acrB* | Antimicrobial activity/Competitive advantage | Same ORF as ARG | 0 |
|  | *acrB* | Antimicrobial activity/Competitive advantage | Same ORF as ARG | 0 |
|  | *acrB* | Antimicrobial activity/Competitive advantage | Same ORF as ARG | 0 |
|  | *acrB* | Antimicrobial activity/Competitive advantage | Same ORF as ARG | 0 |
|  | *msbA* | Immune modulation | Same ORF as ARG | 0 |
|  | *ugd* | Immune modulation | Same ORF as ARG | 0 |
|  | *ugd* | Immune modulation | Same ORF as ARG | 0 |
|  | *ugd* | Immune modulation | Same ORF as ARG | 0 |
| SM | *galU* | Immune modulation | Distinct ORF, separate | 51 |
|  | *adeG* | Biofilm | Same ORF as ARG | 0 |
|  | *adeG* | Biofilm | Same ORF as ARG | 0 |
|  | *ugd* | Immune modulation | Same ORF as ARG | 0 |
|  | *ugd* | Immune modulation | Same ORF as ARG | 0 |
|  | *ugd* | Immune modulation | Same ORF as ARG | 0 |
|  | *ugd* | Immune modulation | Same ORF as ARG | 0 |
| MRH | *adeG* | Biofilm | Same ORF as ARG | 0 |
|  | *adeG* | Biofilm | Same ORF as ARG | 0 |

**Table S16C.** VFDB category and gene counts by site.

| **River** | **Location** | **VF category** | **VF gene** | **VF ORFs** |
| --- | --- | --- | --- | --- |
| Adyar | MM | Antimicrobial activity/Competitive advantage | *acrB* | 11 |
|  |  | Antimicrobial activity/Competitive advantage | *acrA* | 4 |
|  |  | Immune modulation | *ugd* | 3 |
|  |  | Immune modulation | *msbA* | 1 |
|  | SM | Immune modulation | *ugd* | 4 |
|  |  | Biofilm | *adeG* | 2 |
|  |  | Immune modulation | *galU* | 1 |
| Musi | MRH | Biofilm | *adeG* | 2 |

**Table S16D.** Summary of VF ORF–ARG relationships by site.

| **River** | **Location** | **relationship** | **VF ORFs** | **ACCs** |
| --- | --- | --- | --- | --- |
| Adyar | MM | Same ORF as ARG | 19 | 17 |
|  | SM | Distinct ORF, separate | 1 | 1 |
|  |  | Same ORF as ARG | 6 | 6 |
| Musi | MRH | Same ORF as ARG | 2 | 2 |

**Table S17A.** Site prevalence of ACCs with NCyc annotations under primary and relaxed-only thresholds.

| **Threshold** | **River** | **Location** | **Total ACCs** | **ACCs** | **ACCs (%)** | **NCyc ORFs** |
| --- | --- | --- | --- | --- | --- | --- |
| Primary | Adyar | MM | 322 | 5 | 1.55 | 5 |
|  |  | SM | 787 | 3 | 0.38 | 3 |
|  | Musi | MRH | 277 | 1 | 0.36 | 1 |
|  |  | NH | 260 | 0 | 0 | 0 |
|  |  | NRH | 436 | 3 | 0.69 | 3 |
|  |  | SPH | 320 | 0 | 0 | 0 |
| Relaxed-only | Adyar | MM | 322 | 2 | 0.62 | 2 |
|  |  | SM | 787 | 5 | 0.64 | 5 |
|  | Musi | MRH | 277 | 4 | 1.44 | 4 |
|  |  | NH | 260 | 1 | 0.38 | 1 |
|  |  | NRH | 436 | 3 | 0.69 | 3 |
|  |  | SPH | 320 | 1 | 0.31 | 1 |

**Table S17B.** ORF-level NCyc annotations with the closest ARG relationship.

| **Threshold** | **River** | **Location** | **NCyc gene** | **Pathway** | **relationship** | **distance category** | **ARG partners** |
| --- | --- | --- | --- | --- | --- | --- | --- |
| Primary | Adyar | MM | *543728.Vapar_0595* | other_or_unresolved_N_cycle | Distinct ORF, separate | separate_ORFs_between_5_and_10kb | 1 |
|  |  |  | *nasA* | assimilatory_nitrate_reduction | Distinct ORF, separate | same_contig_beyond_10kb | 1 |
|  |  |  | *gdh_K00262* | organic_nitrogen_metabolism | Distinct ORF, separate | separate_ORFs_within_5kb | 1 |
|  |  |  | *ef62402eeb41f049b3f606b486be9dfd* | other_or_unresolved_N_cycle | Distinct ORF, separate | separate_ORFs_between_5_and_10kb | 3 |
|  |  |  | *glnA* | organic_nitrogen_metabolism | Distinct ORF, separate | same_contig_beyond_10kb | 1 |
|  |  | SM | *gs_K00265* | other_or_unresolved_N_cycle | Distinct ORF, separate | separate_ORFs_within_5kb | 1 |
|  |  |  | *glnA* | organic_nitrogen_metabolism | Distinct ORF, separate | separate_ORFs_within_5kb | 1 |
|  |  |  | *glnA* | organic_nitrogen_metabolism | Distinct ORF, separate | separate_ORFs_within_5kb | 1 |
|  | Musi | MRH | *3.84E+08* | other_or_unresolved_N_cycle | Distinct ORF, separate | separate_ORFs_within_5kb | 1 |
|  |  | NRH | *gs_K00266* | other_or_unresolved_N_cycle | Distinct ORF, separate | separate_ORFs_within_5kb | 1 |
|  |  |  | *gs_K00266* | other_or_unresolved_N_cycle | Distinct ORF, separate | separate_ORFs_within_5kb | 1 |
|  |  |  | *gs_K00265* | other_or_unresolved_N_cycle | Distinct ORF, separate | separate_ORFs_within_5kb | 1 |
| Relaxed-only | Adyar | MM | *gs_K00266* | other_or_unresolved_N_cycle | Distinct ORF, separate | separate_ORFs_within_5kb | 1 |
|  |  |  | *gs_K00266* | other_or_unresolved_N_cycle | Distinct ORF, separate | separate_ORFs_within_5kb | 1 |
|  |  | SM | *2.96E+08* | other_or_unresolved_N_cycle | Distinct ORF, separate | separate_ORFs_within_5kb | 1 |
|  |  |  | *gs_K00266* | other_or_unresolved_N_cycle | Distinct ORF, separate | separate_ORFs_within_5kb | 1 |
|  |  |  | *8ebd850849512f6c71b39f9c90e46891* | other_or_unresolved_N_cycle | Distinct ORF, separate | separate_ORFs_within_5kb | 2 |
|  |  |  | *3d983496c6b9ab32d04a473d53161398* | other_or_unresolved_N_cycle | Same ORF as ARG | same_ORF | 1 |
|  |  |  | *bcfa5fca5cbe2e3a88135a94ca50963d* | other_or_unresolved_N_cycle | Same ORF as ARG | same_ORF | 1 |
|  | Musi | MRH | *gs_K00266* | other_or_unresolved_N_cycle | Distinct ORF, separate | separate_ORFs_within_5kb | 1 |
|  |  |  | *hcp* | other_or_unresolved_N_cycle | Distinct ORF, separate | same_contig_beyond_10kb | 1 |
|  |  |  | *1.55E+08* | other_or_unresolved_N_cycle | Distinct ORF, separate | separate_ORFs_within_5kb | 1 |
|  |  |  | *bcfa5fca5cbe2e3a88135a94ca50963d* | other_or_unresolved_N_cycle | Same ORF as ARG | same_ORF | 1 |
|  |  | NH | *gdh_K00262* | organic_nitrogen_metabolism | Distinct ORF, separate | separate_ORFs_within_5kb | 1 |
|  |  | NRH | *gs_K00266* | other_or_unresolved_N_cycle | Distinct ORF, separate | separate_ORFs_within_5kb | 1 |
|  |  |  | *gs_K00266* | other_or_unresolved_N_cycle | Distinct ORF, separate | separate_ORFs_within_5kb | 1 |
|  |  |  | *bcfa5fca5cbe2e3a88135a94ca50963d* | other_or_unresolved_N_cycle | Same ORF as ARG | same_ORF | 1 |
|  |  | SPH | *narB* | denitrification | Distinct ORF, separate | separate_ORFs_within_5kb | 1 |

**Table S17C.** NCyc pathway and gene counts by site.

| **threshold** | **River** | **Location** | **pathway** | **NCyc gene** | **NCyc ORFs** |
| --- | --- | --- | --- | --- | --- |
| Primary | Adyar | MM | assimilatory_nitrate_reduction | *nasA* | 1 |
|  |  |  | organic_nitrogen_metabolism | *gdh_K00262* | 1 |
|  |  |  | organic_nitrogen_metabolism | *glnA* | 1 |
|  |  |  | other_or_unresolved_N_cycle | *543728.Vapar_0595* | 1 |
|  |  |  | other_or_unresolved_N_cycle | *ef62402eeb41f049b3f606b486be9dfd* | 1 |
|  |  | SM | organic_nitrogen_metabolism | *glnA* | 2 |
|  |  |  | other_or_unresolved_N_cycle | *gs_K00265* | 1 |
|  | Musi | MRH | other_or_unresolved_N_cycle | *3.84E+08* | 1 |
|  |  | NRH | other_or_unresolved_N_cycle | *gs_K00266* | 2 |
|  |  |  | other_or_unresolved_N_cycle | *gs_K00265* | 1 |
| Relaxed-only | Adyar | MM | other_or_unresolved_N_cycle | *gs_K00266* | 2 |
|  |  | SM | other_or_unresolved_N_cycle | *2.96E+08* | 1 |
|  |  |  | other_or_unresolved_N_cycle | *3d983496c6b9ab32d04a473d53161398* | 1 |
|  |  |  | other_or_unresolved_N_cycle | *8ebd850849512f6c71b39f9c90e46891* | 1 |
|  |  |  | other_or_unresolved_N_cycle | *bcfa5fca5cbe2e3a88135a94ca50963d* | 1 |
|  |  |  | other_or_unresolved_N_cycle | *gs_K00266* | 1 |
|  | Musi | MRH | other_or_unresolved_N_cycle | *1.55E+08* | 1 |
|  |  |  | other_or_unresolved_N_cycle | *bcfa5fca5cbe2e3a88135a94ca50963d* | 1 |
|  |  |  | other_or_unresolved_N_cycle | *gs_K00266* | 1 |
|  |  |  | other_or_unresolved_N_cycle | *hcp* | 1 |
|  |  | NH | organic_nitrogen_metabolism | *gdh_K00262* | 1 |
|  |  | NRH | other_or_unresolved_N_cycle | *gs_K00266* | 2 |
|  |  |  | other_or_unresolved_N_cycle | *bcfa5fca5cbe2e3a88135a94ca50963d* | 1 |
|  |  | SPH | denitrification | *narB* | 1 |

**Table S17D.** All ARG–NCyc annotation pairs-level genomic coordinates and distances.

| **Threshold** | **River** | **Location** | **NCyc gene** | **Pathway** | **Relationship** | **Distance (bp)** |
| --- | --- | --- | --- | --- | --- | --- |
| Primary | Adyar | MM | *gdh_K00262* | organic_nitrogen_metabolism | Distinct ORF, separate | 3778 |
|  |  |  | *ef62402eeb41f049b3f606b486be9dfd* | other_or_unresolved_N_cycle | Distinct ORF, separate | 5321 |
|  |  |  | *ef62402eeb41f049b3f606b486be9dfd* | other_or_unresolved_N_cycle | Distinct ORF, separate | 6835 |
|  |  |  | *543728.Vapar_0595* | other_or_unresolved_N_cycle | Distinct ORF, separate | 8776 |
|  |  |  | *ef62402eeb41f049b3f606b486be9dfd* | other_or_unresolved_N_cycle | Distinct ORF, separate | 10011 |
|  |  |  | *nasA* | assimilatory_nitrate_reduction | Distinct ORF, separate | 33854 |
|  |  |  | *glnA* | organic_nitrogen_metabolism | Distinct ORF, separate | 38487 |
|  |  | SM | *glnA* | organic_nitrogen_metabolism | Distinct ORF, separate | 224 |
|  |  |  | *glnA* | organic_nitrogen_metabolism | Distinct ORF, separate | 224 |
|  |  |  | *gs_K00265* | other_or_unresolved_N_cycle | Distinct ORF, separate | 2320 |
|  | Musi | MRH | *3.84E+08* | other_or_unresolved_N_cycle | Distinct ORF, separate | 2792 |
|  |  | NRH | *gs_K00266* | other_or_unresolved_N_cycle | Distinct ORF, separate | 160 |
|  |  |  | *gs_K00266* | other_or_unresolved_N_cycle | Distinct ORF, separate | 161 |
|  |  |  | *gs_K00265* | other_or_unresolved_N_cycle | Distinct ORF, separate | 2437 |
| Relaxed-only | Adyar | MM | *gs_K00266* | other_or_unresolved_N_cycle | Distinct ORF, separate | 146 |
|  |  |  | *gs_K00266* | other_or_unresolved_N_cycle | Distinct ORF, separate | 151 |
|  |  | SM | *2.96E+08* | other_or_unresolved_N_cycle | Distinct ORF, separate | 11 |
|  |  |  | *gs_K00266* | other_or_unresolved_N_cycle | Distinct ORF, separate | 158 |
|  |  |  | *8ebd850849512f6c71b39f9c90e46891* | other_or_unresolved_N_cycle | Distinct ORF, separate | 2369 |
|  |  |  | *8ebd850849512f6c71b39f9c90e46891* | other_or_unresolved_N_cycle | Distinct ORF, separate | 3563 |
|  |  |  | *3d983496c6b9ab32d04a473d53161398* | other_or_unresolved_N_cycle | Same ORF as ARG | 0 |
|  |  |  | *bcfa5fca5cbe2e3a88135a94ca50963d* | other_or_unresolved_N_cycle | Same ORF as ARG | 0 |
|  | Musi | MRH | *1.55E+08* | other_or_unresolved_N_cycle | Distinct ORF, separate | 11 |
|  |  |  | *gs_K00266* | other_or_unresolved_N_cycle | Distinct ORF, separate | 139 |
|  |  |  | *hcp* | other_or_unresolved_N_cycle | Distinct ORF, separate | 19850 |
|  |  |  | *bcfa5fca5cbe2e3a88135a94ca50963d* | other_or_unresolved_N_cycle | Same ORF as ARG | 0 |
|  |  | NH | *gdh_K00262* | organic_nitrogen_metabolism | Distinct ORF, separate | 4576 |
|  |  | NRH | *gs_K00266* | other_or_unresolved_N_cycle | Distinct ORF, separate | 151 |
|  |  |  | *gs_K00266* | other_or_unresolved_N_cycle | Distinct ORF, separate | 151 |
|  |  |  | *bcfa5fca5cbe2e3a88135a94ca50963d* | other_or_unresolved_N_cycle | Same ORF as ARG | 0 |
|  |  | SPH | *narB* | denitrification | Distinct ORF, separate | 262 |

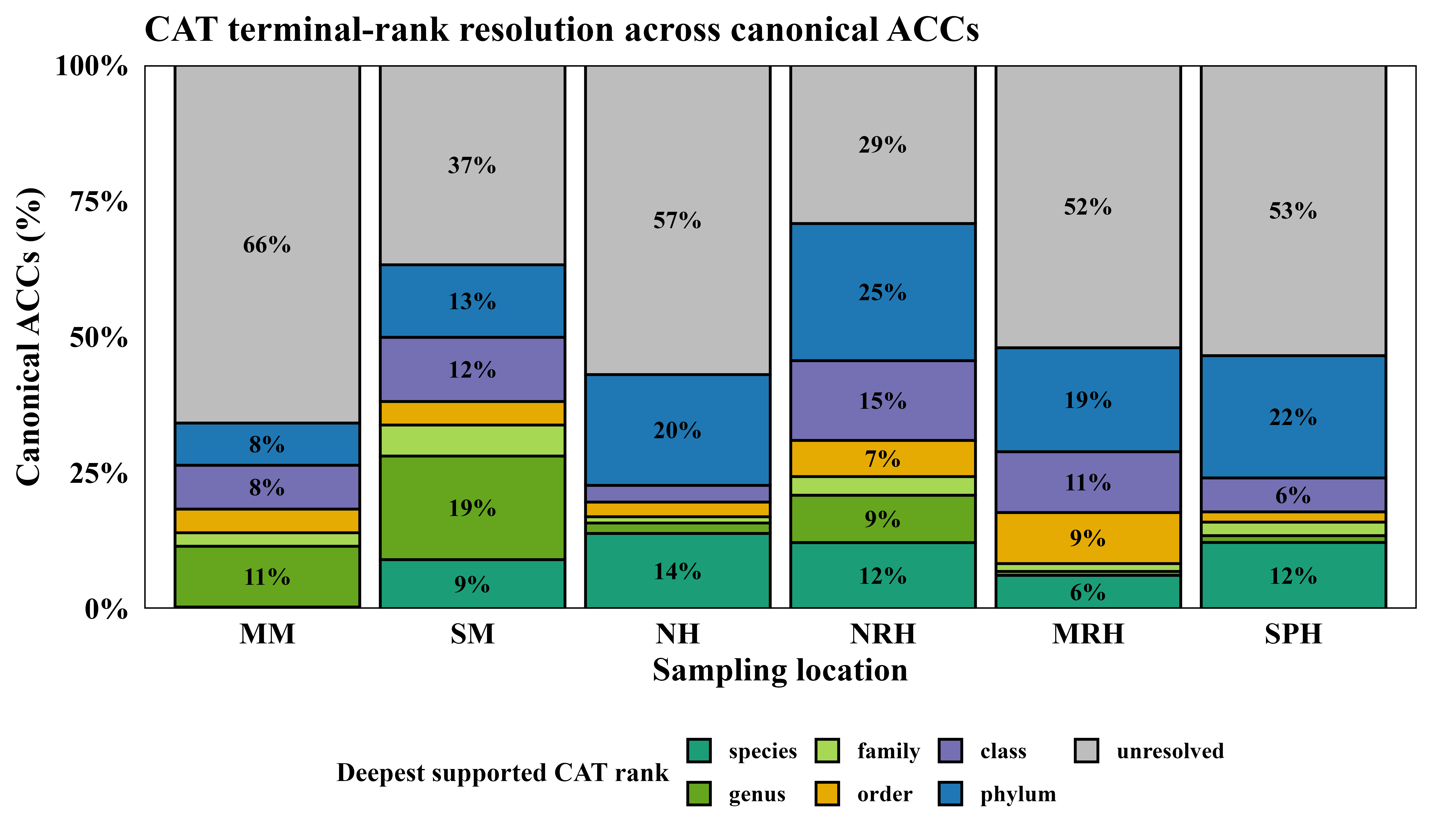
**Figure S6.** Taxonomic rank resolution of canonical ACCs across river sampling locations.

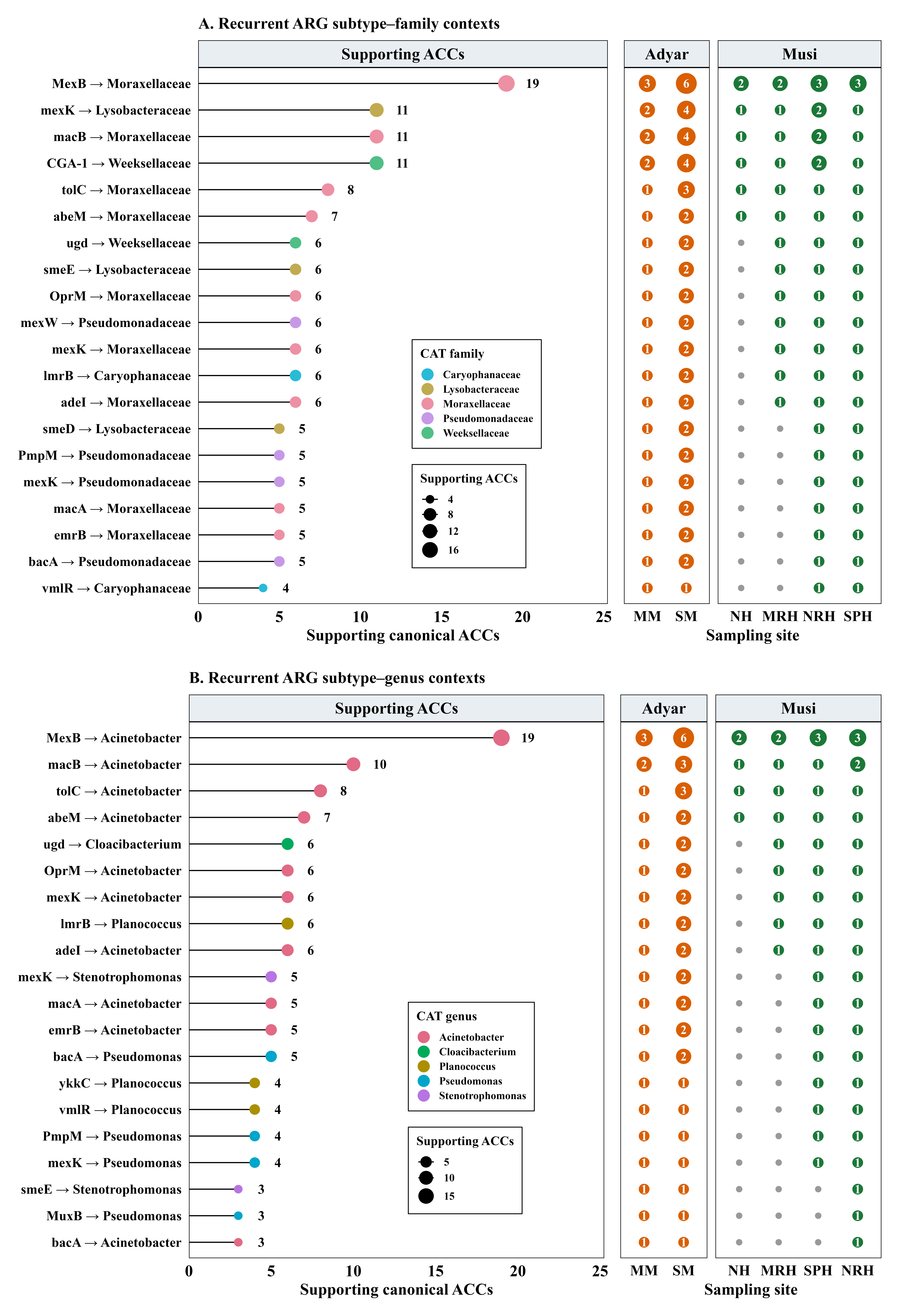

**Figure S7.** Recurrent ARG subtype **(A)** family and **(B)** genus taxonomic contexts across riverine sampling sites.

**Table S18A.** Deepest supported CAT rank of canonical ACCs by site (ACC counts, with the percentage of ACCs resolved and unresolved).

| **River** | **Site** | **Canonical ACCs (n)** | **Species** | **Genus** | **Family** | **Order** | **Class** | **Phylum** | **Unresolved** | **Resolved to at least phylum (n)** | **Resolved (%)** | **Unresolved (%)** |
| --- | --- | --- | --- | --- | --- | --- | --- | --- | --- | --- | --- | --- |
| Adyar | MM | 322 | 1 | 36 | 8 | 14 | 26 | 25 | 212 | 110 | 34.2 | 65.8 |
|  | SM | 787 | 71 | 150 | 45 | 34 | 93 | 105 | 289 | 498 | 63.3 | 36.7 |
| Musi | NH | 260 | 36 | 5 | 3 | 7 | 8 | 53 | 148 | 112 | 43.1 | 56.9 |
|  | NRH | 436 | 53 | 38 | 15 | 29 | 64 | 110 | 127 | 309 | 70.9 | 29.1 |
|  | MRH | 277 | 17 | 2 | 4 | 26 | 31 | 53 | 144 | 133 | 48 | 52 |
|  | SPH | 320 | 39 | 4 | 8 | 6 | 20 | 72 | 171 | 149 | 46.6 | 53.4 |
| Total | All sites | 2402 | 217 | 235 | 83 | 116 | 242 | 418 | 1091 | 1311 | 54.6 | 45.4 |

**Table S18B.** Recurrent ARG subtype phylum associations supported by at least two ACCs.

| **ARG class** | **ARG subtype** | **CAT phylum** | **Supporting ACCs** | **Supporting sites** |
| --- | --- | --- | --- | --- |
| multidrug | *MexB* | *Pseudomonadota* | 75 | MM, MRH, NH, NRH, SM, SPH |
| multidrug | *mexK* | *Pseudomonadota* | 73 | MM, MRH, NH, NRH, SM, SPH |
| bacitracin | *bacA* | *Pseudomonadota* | 70 | MM, MRH, NH, NRH, SM, SPH |
| multidrug | *smeE* | *Pseudomonadota* | 63 | MM, MRH, NH, NRH, SM, SPH |
| novobiocin | *novA* | *Pseudomonadota* | 50 | MM, MRH, NH, NRH, SM, SPH |
| multidrug | *MuxB* | *Pseudomonadota* | 36 | MM, MRH, NRH, SM |
| multidrug | *mexW* | *Pseudomonadota* | 33 | MRH, NH, NRH, SM, SPH |
| polymyxin | *rosB* | *Pseudomonadota* | 33 | MRH, NRH, SM, SPH |
| multidrug | *MexF* | *Pseudomonadota* | 32 | MRH, NH, NRH, SM, SPH |
| multidrug | *msbA* | *Pseudomonadota* | 28 | MM, MRH, NH, NRH, SM |
| novobiocin | *novA* | *Actinomycetota* | 27 | NH, NRH, SPH |
| multidrug | *OprM* | *Pseudomonadota* | 23 | MM, NH, SM |
| macrolide-lincosamide-streptogramin | *macB* | *Pseudomonadota* | 23 | MM, MRH, NRH, SM, SPH |
| multidrug | *oqxB* | *Pseudomonadota* | 22 | MM, MRH, NH, NRH, SM, SPH |
| polymyxin | *ugd* | *Pseudomonadota* | 20 | MM, MRH, SM |
| multidrug | *ceoB* | *Pseudomonadota* | 19 | MM, MRH, NH, NRH, SM, SPH |
| macrolide-lincosamide-streptogramin | *ole(C)* | *Actinomycetota* | 16 | NRH, SM, SPH |
| multidrug | *abeM* | *Pseudomonadota* | 15 | MM, NRH, SM |
| multidrug | *acrF* | *Pseudomonadota* | 15 | MM, NH, NRH, SM, SPH |
| multidrug | *adeI* | *Pseudomonadota* | 15 | MM, SM |
| multidrug | *MexD* | *Pseudomonadota* | 14 | MM, NH, NRH, SM, SPH |
| multidrug | *RanA* | *Bacteroidota* | 14 | NRH, SM |
| multidrug | *MuxC* | *Pseudomonadota* | 12 | MM, NRH, SM |
| polymyxin | *rosA* | *Pseudomonadota* | 12 | MM, MRH, NRH, SM, SPH |
| multidrug | *smeD* | *Pseudomonadota* | 12 | MM, MRH, NH, NRH, SM |
| polymyxin | *ugd* | *Bacteroidota* | 12 | MM, SM |
| beta lactam | *CGA-1* | *Bacteroidota* | 11 | MM, SM |
| tetracycline | *tetA(48)* | *Actinomycetota* | 11 | MRH, NRH, SPH |
| multidrug | *tolC* | *Pseudomonadota* | 11 | MM, SM |
| mupirocin | *Bifidobacteria intrinsic ileS conferring resistance to mupirocin* | *Actinomycetota* | 9 | MRH, NRH, SPH |
| multidrug | *abeS* | *Pseudomonadota* | 9 | MM, MRH, NRH, SM |
| bacitracin | *bacA* | *Bacillota* | 9 | MRH, NRH, SM |
| multidrug | *qacE* | *Pseudomonadota* | 9 | NRH, SM |
| multidrug | *emrB* | *Pseudomonadota* | 8 | MM, NRH, SM |
| sulfonamide | *sul4* | *Chloroflexota* | 8 | NH, NRH, SM |
| multidrug | *MexA* | *Pseudomonadota* | 7 | SM |
| multidrug | *bpeF* | *Pseudomonadota* | 7 | MRH, NH, NRH, SM, SPH |
| macrolide-lincosamide-streptogramin | *macA* | *Pseudomonadota* | 7 | NH, SM |
| novobiocin | *novA* | *Acidobacteriota* | 7 | NH, NRH, SPH |
| novobiocin | *novA* | *Cyanobacteriota* | 7 | MRH, NRH |
| multidrug | *PmpM* | *Pseudomonadota* | 6 | SM |
| multidrug | *amrB* | *Pseudomonadota* | 6 | MRH, SM, SPH |
| macrolide-lincosamide-streptogramin | *lmrB* | *Bacillota* | 6 | NRH |
| multidrug | *mdtF* | *Pseudomonadota* | 6 | MRH, NRH, SM |
| multidrug | *msbA* | *Bacteroidota* | 6 | MRH, NH, NRH, SM |
| puromycin | *puromycin resistance protein* | *Actinomycetota* | 6 | SPH |
| multidrug | *Escherichia coli mdfA* | *Pseudomonadota* | 5 | MM, SM |
| multidrug | *MuxB* | *Acidobacteriota* | 5 | NH, NRH, SPH |
| multidrug | *OprN* | *Pseudomonadota* | 5 | MRH, NRH, SM, SPH |
| multidrug | *RanB* | *Bacteroidota* | 5 | SM |
| rifamycin | *RbpA* | *Actinomycetota* | 5 | SPH |
| bleomycin | *bleomycin resistance protein* | *Pseudomonadota* | 5 | NRH |
| multidrug | *efrB* | *Bacillota* | 5 | MM, NH, NRH, SM |
| macrolide-lincosamide-streptogramin | *lnu(H)* | *Bacteroidota* | 5 | SM |
| multidrug | *mdtE* | *Pseudomonadota* | 5 | MM, SM |
| multidrug | *mexN* | *Pseudomonadota* | 5 | NRH, SM |
| novobiocin | *novA* | *Chloroflexota* | 5 | MRH, NH, SM |
| multidrug | *qacH* | *Pseudomonadota* | 5 | MM, MRH, NRH, SM |
| macrolide-lincosamide-streptogramin | *vat(F)* | *Cyanobacteriota* | 5 | MRH, NRH |
| fosfomycin | *Acinetobacter baumannii AbaF* | *Actinomycetota* | 4 | SPH |
| multidrug | *Escherichia coli emrE* | *Pseudomonadota* | 4 | NH, NRH, SM |
| multidrug | *MexB* | *Campylobacterota* | 4 | SM |
| multidrug | *MexE* | *Pseudomonadota* | 4 | MRH, SM, SPH |
| multidrug | *Pseudomonas aeruginosa emrE* | *Pseudomonadota* | 4 | NRH, SM |
| multidrug | *adeF* | *Pseudomonadota* | 4 | MRH, NRH, SM, SPH |
| trimethoprim | *dfrA3* | *Pseudomonadota* | 4 | NRH, SM, SPH |
| multidrug | *efrA* | *Actinomycetota* | 4 | NH, NRH, SM, SPH |
| multidrug | *efrA* | *Bacillota* | 4 | NH, SM |
| other_peptide_antibiotics | *microcin efflux pumu gene yojI* | *Pseudomonadota* | 4 | MM, SM |
| novobiocin | *novA* | *Bacillota* | 4 | NH, SM, SPH |
| multidrug | *smeF* | *Pseudomonadota* | 4 | NRH, SM |
| macrolide-lincosamide-streptogramin | *vmlR* | *Bacillota* | 4 | NRH |
| multidrug | *ykkC* | *Bacillota* | 4 | NRH |
| aminoglycoside | *AAC(3)-Id* | *Pseudomonadota* | 3 | MRH, NRH |
| multidrug | *MexB* | *Acidobacteriota* | 3 | NH |
| multidrug | *MexC* | *Pseudomonadota* | 3 | MM, SM |
| multidrug | *MexF* | *Planctomycetota* | 3 | NRH |
| multidrug | *acrF* | *Thermodesulfobacteriota* | 3 | NH, SM |
| polymyxin | *arnA* | *Pseudomonadota* | 3 | MM, NH, NRH |
| multidrug | *emrA* | *Pseudomonadota* | 3 | MM, MRH, NRH |
| macrolide-lincosamide-streptogramin | *lsa(B)* | *Bacillota* | 3 | NH |
| macrolide-lincosamide-streptogramin | *macB* | *Cyanobacteriota* | 3 | MRH, NRH |
| macrolide-lincosamide-streptogramin | *macB* | *Acidobacteriota* | 3 | NH |
| multidrug | *mdsB* | *Pseudomonadota* | 3 | MRH, NRH |
| multidrug | *mdtH* | *Pseudomonadota* | 3 | MM, SM |
| novobiocin | *novA* | *Gemmatimonadota* | 3 | NRH, SPH |
| novobiocin | *novA* | *Synergistota* | 3 | NH, SM |
| multidrug | *sdeY* | *Pseudomonadota* | 3 | SM |
| sulfonamide | *sul1* | *Pseudomonadota* | 3 | MRH, SM |
| tetracycline | *tetA(48)* | *Pseudomonadota* | 3 | MRH, NRH |
| aminoglycoside | *AAC(6')-Ib8* | *Bacteroidota* | 2 | SM |
| aminoglycoside | *AAC(6')-Isa* | *Actinomycetota* | 2 | SPH |
| beta lactam | *CRD3-1* | *Pseudomonadota* | 2 | SPH |
| fosfomycin | *FosX* | *Pseudomonadota* | 2 | MRH, SM |
| fosfomycin | *FosX* | *Cyanobacteriota* | 2 | MRH |
| beta lactam | *Klebsiella pneumoniae OmpK37* | *Pseudomonadota* | 2 | MM |
| beta lactam | *LCR-1* | *Pseudomonadota* | 2 | NH, SM |
| multidrug | *MexB* | *Thermodesulfobacteriota* | 2 | NH, SM |
| multidrug | *MexE* | *Acidobacteriota* | 2 | NH |
| multidrug | *MuxC* | *Acidobacteriota* | 2 | NRH, SPH |
| beta lactam | *OXA-427* | *Pseudomonadota* | 2 | SM |
| beta lactam | *OXA-464* | *Campylobacterota* | 2 | SM |
| multidrug | *OpmH* | *Pseudomonadota* | 2 | SM |
| multidrug | *OprM* | *Acidobacteriota* | 2 | NH, SM |
| aminoglycoside | *aadS* | *Bacteroidota* | 2 | NH, SM |
| multidrug | *abeS* | *Methanobacteriota* | 2 | NH, SM |
| multidrug | *acrF* | *Acidobacteriota* | 2 | NH, SPH |
| bacitracin | *bacA* | *Thermodesulfobacteriota* | 2 | NH, SM |
| multidrug | *ceoB* | *Campylobacterota* | 2 | SM |
| multidrug | *efrA* | *Chloroflexota* | 2 | SM |
| multidrug | *efrB* | *Chloroflexota* | 2 | NH |
| multidrug | *emrD* | *Pseudomonadota* | 2 | MM, SM |
| macrolide-lincosamide-streptogramin | *ere(D)* | *Bacteroidota* | 2 | SM |
| macrolide-lincosamide-streptogramin | *erm(37)* | *Actinomycetota* | 2 | SPH |
| macrolide-lincosamide-streptogramin | *lmrD* | *Chloroflexota* | 2 | NH, NRH |
| multidrug | *mdtF* | *Acidobacteriota* | 2 | NH |
| macrolide-lincosamide-streptogramin | *mel* | *Bacillota* | 2 | NH |
| multidrug | *mexQ* | *Acidobacteriota* | 2 | MRH |
| multidrug | *mexV* | *Pseudomonadota* | 2 | SM |
| macrolide-lincosamide-streptogramin | *mph(A)* | *Myxococcota* | 2 | NH, SM |
| multidrug | *msbA* | *Acidobacteriota* | 2 | NRH, SPH |
| pleuromutilin_tiamulin | *msr(G)* | *Bacillota* | 2 | MM, NRH |
| multidrug | *mtrD* | *Pseudomonadota* | 2 | MM, NRH |
| mupirocin | *mupB* | *Chloroflexota* | 2 | NRH, SM |
| tetracycline | *otr(A)* | *Actinomycetota* | 2 | SPH |
| polymyxin | *pmrF* | *Pseudomonadota* | 2 | MM |
| quinolone | *qepA* | *Actinomycetota* | 2 | SPH |
| polymyxin | *rosA* | *Thermodesulfobacteriota* | 2 | NH, SM |
| polymyxin | *rosB* | *Acidobacteriota* | 2 | NH, SM |
| rifamycin | *rphA* | *Actinomycetota* | 2 | SPH |
| multidrug | *smeE* | *Gemmatimonadota* | 2 | NH |
| sulfonamide | *sul2* | *Pseudomonadota* | 2 | NRH |
| tetracycline | *tet(34)* | *Pseudomonadota* | 2 | SM |
| tetracycline | *tet(Q)* | *Bacteroidota* | 2 | NH, SM |
| tetracycline | *tetA(48)* | *Bacillota* | 2 | NRH |
| tetracycline | *tetB(46)* | *Actinomycetota* | 2 | MRH, SPH |
| tetracycline | *tetB(46)* | *Bacillota* | 2 | NRH |

**Table S18C.** Recurrent ARG subtype family associations supported by at least two ACCs.

| **ARG class** | **ARG subtype** | **CAT family** | **Supporting ACCs (n)** | **Supporting sites** |
| --- | --- | --- | --- | --- |
| multidrug | *MexB* | *Moraxellaceae* | 19 | MM, SM |
| beta lactam | *CGA-1* | *Weeksellaceae* | 11 | MM, SM |
| macrolide-lincosamide-streptogramin | *macB* | *Moraxellaceae* | 11 | MM, NRH, SM |
| multidrug | *mexK* | *Lysobacteraceae* | 11 | MRH, NRH, SM |
| multidrug | *tolC* | *Moraxellaceae* | 8 | MM, SM |
| multidrug | *abeM* | *Moraxellaceae* | 7 | MM, SM |
| multidrug | *OprM* | *Moraxellaceae* | 6 | MM, SM |
| multidrug | *adeI* | *Moraxellaceae* | 6 | MM, SM |
| macrolide-lincosamide-streptogramin | *lmrB* | *Caryophanaceae* | 6 | NRH |
| multidrug | *mexK* | *Moraxellaceae* | 6 | MM, SM |
| multidrug | *mexW* | *Pseudomonadaceae* | 6 | SM |
| multidrug | *smeE* | *Lysobacteraceae* | 6 | MM, SM |
| polymyxin | *ugd* | *Weeksellaceae* | 6 | SM |
| multidrug | *PmpM* | *Pseudomonadaceae* | 5 | SM |
| bacitracin | *bacA* | *Pseudomonadaceae* | 5 | SM |
| multidrug | *emrB* | *Moraxellaceae* | 5 | SM |
| macrolide-lincosamide-streptogramin | *macA* | *Moraxellaceae* | 5 | SM |
| multidrug | *mexK* | *Pseudomonadaceae* | 5 | SM |
| multidrug | *smeD* | *Lysobacteraceae* | 5 | MM, SM |
| multidrug | *MuxB* | *Pseudomonadaceae* | 4 | SM |
| multidrug | *oqxB* | *Comamonadaceae* | 4 | NRH |
| macrolide-lincosamide-streptogramin | *vmlR* | *Caryophanaceae* | 4 | NRH |
| multidrug | *ykkC* | *Caryophanaceae* | 4 | NRH |
| multidrug | *MexB* | *Burkholderiaceae* | 3 | MM, NRH |
| multidrug | *MexB* | *Arcobacteraceae* | 3 | SM |
| multidrug | *MexB* | *Pseudomonadaceae* | 3 | SM |
| multidrug | *MuxC* | *Pseudomonadaceae* | 3 | SM |
| bacitracin | *bacA* | *Moraxellaceae* | 3 | MM, SM |
| bacitracin | *bacA* | *Paracoccaceae* | 3 | MM, NRH |
| novobiocin | *novA* | *Paracoccaceae* | 3 | MM, MRH, SM |
| novobiocin | *novA* | *Synergistaceae* | 3 | NH, SM |
| multidrug | *smeE* | *Burkholderiaceae* | 3 | NRH, SPH |
| multidrug | *smeE* | *Alcaligenaceae* | 3 | SM |
| sulfonamide | *sul4* | *Anaerolineaceae* | 3 | NH, SM |
| multidrug | *Escherichia coli mdfA* | *Moraxellaceae* | 2 | MM, SM |
| multidrug | *MexB* | *Alcaligenaceae* | 2 | SM |
| multidrug | *MexB* | *Alteromonadaceae* | 2 | SM |
| multidrug | *MuxB* | *Comamonadaceae* | 2 | NRH, SM |
| multidrug | *MuxB* | *Alcaligenaceae* | 2 | SM |
| beta lactam | *OXA-427* | *Aeromonadaceae* | 2 | SM |
| beta lactam | *OXA-464* | *Arcobacteraceae* | 2 | SM |
| multidrug | *OpmH* | *Pseudomonadaceae* | 2 | SM |
| multidrug | *Pseudomonas aeruginosa emrE* | *Pseudomonadaceae* | 2 | SM |
| multidrug | *RanB* | *Weeksellaceae* | 2 | SM |
| multidrug | *acrF* | *Desulfobulbaceae* | 2 | NH, SM |
| multidrug | *amrB* | *Rhodocyclaceae* | 2 | SM |
| bacitracin | *bacA* | *Desulfobulbaceae* | 2 | NH, SM |
| bacitracin | *bacA* | *Caryophanaceae* | 2 | NRH |
| bacitracin | *bacA* | *Filifactoraceae* | 2 | SM |
| multidrug | *ceoB* | *Arcobacteraceae* | 2 | SM |
| macrolide-lincosamide-streptogramin | *lnu(H)* | *Flavobacteriaceae* | 2 | SM |
| macrolide-lincosamide-streptogramin | *macB* | *Aeromonadaceae* | 2 | SM |
| multidrug | *mexK* | *Comamonadaceae* | 2 | SM |
| multidrug | *mexW* | *Burkholderiaceae* | 2 | NRH |
| multidrug | *msbA* | *Aeromonadaceae* | 2 | SM |
| novobiocin | *novA* | *Anaerolineaceae* | 2 | NH, SM |
| novobiocin | *novA* | *Lysobacteraceae* | 2 | NRH, SM |
| novobiocin | *novA* | *Solirubrobacteraceae* | 2 | NRH, SPH |
| novobiocin | *novA* | *Rhizobiaceae* | 2 | NRH |
| polymyxin | *rosA* | *Desulfobulbaceae* | 2 | NH, SM |
| polymyxin | *rosA* | *Paracoccaceae* | 2 | MM, NRH |
| multidrug | *smeE* | *Paracoccaceae* | 2 | MM, NRH |
| multidrug | *smeE* | *Moraxellaceae* | 2 | SM |
| multidrug | *smeF* | *Lysobacteraceae* | 2 | SM |
| polymyxin | *ugd* | *Lysobacteraceae* | 2 | SM |
| polymyxin | *ugd* | *Paracoccaceae* | 2 | MM |

**Table S18D.** Recurrent ARG subtype genus associations supported by at least two ACCs.

| **ARG class** | **ARG subtype** | **CAT genus** | **Supporting ACCs** | **Supporting sites** |
| --- | --- | --- | --- | --- |
| multidrug | *MexB* | *Acinetobacter* | 19 | MM, SM |
| macrolide-lincosamide-streptogramin | *macB* | *Acinetobacter* | 10 | MM, SM |
| multidrug | *tolC* | *Acinetobacter* | 8 | MM, SM |
| multidrug | *abeM* | *Acinetobacter* | 7 | MM, SM |
| multidrug | *OprM* | *Acinetobacter* | 6 | MM, SM |
| multidrug | *adeI* | *Acinetobacter* | 6 | MM, SM |
| macrolide-lincosamide-streptogramin | *lmrB* | *Planococcus* | 6 | NRH |
| multidrug | *mexK* | *Acinetobacter* | 6 | MM, SM |
| polymyxin | *ugd* | *Cloacibacterium* | 6 | SM |
| bacitracin | *bacA* | *Pseudomonas* | 5 | SM |
| multidrug | *emrB* | *Acinetobacter* | 5 | SM |
| macrolide-lincosamide-streptogramin | *macA* | *Acinetobacter* | 5 | SM |
| multidrug | *mexK* | *Stenotrophomonas* | 5 | SM |
| multidrug | *PmpM* | *Pseudomonas* | 4 | SM |
| multidrug | *mexK* | *Pseudomonas* | 4 | SM |
| macrolide-lincosamide-streptogramin | *vmlR* | *Planococcus* | 4 | NRH |
| multidrug | *ykkC* | *Planococcus* | 4 | NRH |
| multidrug | *MuxB* | *Pseudomonas* | 3 | SM |
| multidrug | *MuxC* | *Pseudomonas* | 3 | SM |
| bacitracin | *bacA* | *Acinetobacter* | 3 | MM, SM |
| multidrug | *oqxB* | *Comamonas* | 3 | NRH |
| multidrug | *smeD* | *Arenimonas* | 3 | SM |
| multidrug | *smeE* | *Stenotrophomonas* | 3 | MM, SM |
| multidrug | *smeE* | *Arenimonas* | 3 | SM |
| beta lactam | *CGA-1* | *Cloacibacterium* | 2 | MM, SM |
| multidrug | *Escherichia coli mdfA* | *Acinetobacter* | 2 | MM, SM |
| multidrug | *MexB* | *Alishewanella* | 2 | SM |
| multidrug | *MexB* | *Pseudomonas* | 2 | SM |
| beta lactam | *OXA-427* | *Aeromonas* | 2 | SM |
| multidrug | *Pseudomonas aeruginosa emrE* | *Pseudomonas* | 2 | SM |
| multidrug | *RanB* | *Cloacibacterium* | 2 | SM |
| bacitracin | *bacA* | *Acetoanaerobium* | 2 | SM |
| bacitracin | *bacA* | *Paracoccus* | 2 | MM |
| bacitracin | *bacA* | *Planococcus* | 2 | NRH |
| macrolide-lincosamide-streptogramin | *macB* | *Aeromonas* | 2 | SM |
| multidrug | *mexK* | *Arenimonas* | 2 | SM |
| multidrug | *mexW* | *Geopseudomonas* | 2 | SM |
| multidrug | *mexW* | *Pseudomonas* | 2 | SM |
| multidrug | *msbA* | *Aeromonas* | 2 | SM |
| polymyxin | *rosA* | *Desulfobulbus* | 2 | NH, SM |
| multidrug | *smeD* | *Stenotrophomonas* | 2 | MM, SM |
| multidrug | *smeE* | *Acinetobacter* | 2 | SM |
| multidrug | *smeF* | *Stenotrophomonas* | 2 | SM |
| tetracycline | *tetB(46)* | *Exiguobacterium** | 2 | NRH |
| polymyxin | *ugd* | *Paracoccus* | 2 | MM |
| polymyxin | *ugd* | *Stenotrophomonas* | 2 | SM |

**Table S18E.** Phylum-level composition of CAT-assigned ACCs by site (ACC counts).

| **CAT phylum** | **MM** | **SM** | **NH** | **NRH** | **MRH** | **SPH** | **All sites** |
| --- | --- | --- | --- | --- | --- | --- | --- |
| *Pseudomonadota* | 102 | 388 | 40 | 220 | 101 | 48 | 899 |
| *Actinomycetota* | 1 | 4 | 4 | 20 | 8 | 75 | 112 |
| *Bacteroidota* | 2 | 54 | 7 | 3 | 1 | 0 | 67 |
| *Bacillota* | 5 | 17 | 9 | 29 | 1 | 1 | 62 |
| *Acidobacteriota* | 0 | 2 | 17 | 8 | 2 | 12 | 41 |
| *Chloroflexota* | 0 | 9 | 12 | 9 | 2 | 0 | 32 |
| *Cyanobacteriota* | 0 | 0 | 0 | 7 | 12 | 0 | 19 |
| *Thermodesulfobacteriota* | 0 | 7 | 8 | 0 | 0 | 0 | 15 |
| *Gemmatimonadota* | 0 | 0 | 2 | 2 | 1 | 7 | 12 |
| *Campylobacterota* | 0 | 10 | 0 | 0 | 0 | 0 | 10 |
| *Myxococcota* | 0 | 1 | 2 | 2 | 0 | 2 | 7 |
| *Nitrospirota* | 0 | 0 | 0 | 0 | 3 | 3 | 6 |
| *Candidatus Aminicenantota* | 0 | 1 | 3 | 0 | 0 | 0 | 4 |
| *Methanobacteriota* | 0 | 1 | 3 | 0 | 0 | 0 | 4 |
| *Planctomycetota* | 0 | 0 | 0 | 4 | 0 | 0 | 4 |
| *Synergistota* | 0 | 2 | 1 | 0 | 0 | 0 | 3 |
| *Verrucomicrobiota* | 0 | 0 | 1 | 1 | 1 | 0 | 3 |
| *Candidatus Cryosericota* | 0 | 1 | 1 | 0 | 0 | 0 | 2 |
| *Armatimonadota* | 0 | 0 | 0 | 1 | 0 | 0 | 1 |
| *Candidatus Binatota* | 0 | 0 | 0 | 1 | 0 | 0 | 1 |
| *Candidatus Hydrogenedentota* | 0 | 1 | 0 | 0 | 0 | 0 | 1 |
| *Candidatus Moraniibacteriota* | 0 | 0 | 1 | 0 | 0 | 0 | 1 |
| *Candidatus Tectimicrobiota* | 0 | 0 | 0 | 0 | 0 | 1 | 1 |
| *Deinococcota* | 0 | 0 | 0 | 0 | 1 | 0 | 1 |
| *Thermoplasmatota* | 0 | 0 | 1 | 0 | 0 | 0 | 1 |

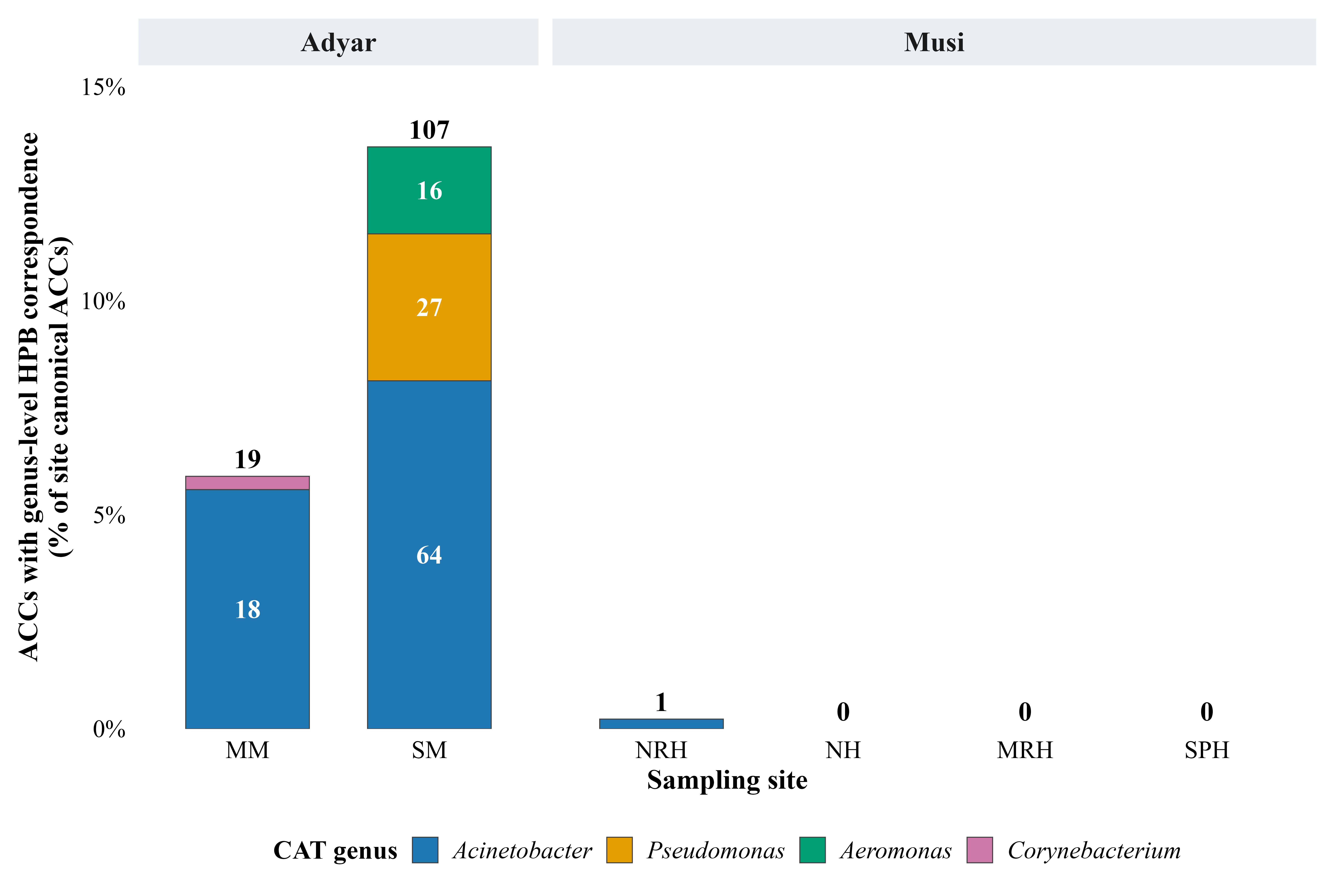
**Figure S8.** Genus composition of canonical ACCs with genus-level correspondence to the HPB reference list, by site. Bar height is the percentage of canonical ACCs at each site; numbers are ACC counts.

**Table S19A.** Distribution of ACCs among evidence classes C1-C5 by site (n and % of canonical ACCs per site).

| **Evidence class** | **MM** | **SM** | **NH** | **NRH** | **MRH** | **SPH** | **All sites** |
| --- | --- | --- | --- | --- | --- | --- | --- |
| Canonical ACCs (n) | 322 | 787 | 260 | 436 | 277 | 320 | 2402 |
| C1: species-confirmed HPB + mobility evidence | 0 (0.00%) | 0 (0.00%) | 0 (0.00%) | 0 (0.00%) | 0 (0.00%) | 0 (0.00%) | 0 (0.00%) |
| C2: species-confirmed HPB, no mobility evidence | 0 (0.00%) | 0 (0.00%) | 0 (0.00%) | 0 (0.00%) | 0 (0.00%) | 0 (0.00%) | 0 (0.00%) |
| C3: mobility evidence, no HPB correspondence | 17 (5.28%) | 31 (3.94%) | 14 (5.38%) | 10 (2.29%) | 1 (0.36%) | 0 (0.00%) | 73 (3.04%) |
| C3a: genus-level HPB correspondence + mobility evidence | 1 (0.31%) | 1 (0.13%) | 0 (0.00%) | 0 (0.00%) | 0 (0.00%) | 0 (0.00%) | 2 (0.08%) |
| C4: genus-level HPB correspondence, no mobility evidence | 18 (5.59%) | 106 (13.47%) | 0 (0.00%) | 1 (0.23%) | 0 (0.00%) | 0 (0.00%) | 125 (5.20%) |
| C5: neither HPB correspondence nor mobility evidence | 286 (88.82%) | 649 (82.47%) | 246 (94.62%) | 425 (97.48%) | 276 (99.64%) | 320 (100.00%) | 2202 (91.67%) |

**Table S19B.** ACCs with genus-level HPB-list correspondence, by CAT-assigned genus and site.

| **CAT genus** | **MM** | **SM** | **NH** | **NRH** | **MRH** | **SPH** | **Total** | **% of HPB-genus ACCs** |
| --- | --- | --- | --- | --- | --- | --- | --- | --- |
| *Acinetobacter* | 18 | 64 | 0 | 1 | 0 | 0 | 83 | 65.4 |
| *Pseudomonas* | 0 | 27 | 0 | 0 | 0 | 0 | 27 | 21.3 |
| *Aeromonas* | 0 | 16 | 0 | 0 | 0 | 0 | 16 | 12.6 |
| *Corynebacterium* | 1 | 0 | 0 | 0 | 0 | 0 | 1 | 0.8 |
| All | 19 | 107 | 0 | 1 | 0 | 0 | 127 | 100 |

**Table S19C.** ACCs combining genus-level HPB-list correspondence with mobility evidence (class C3a): contig characteristics, taxonomic support and mobility evidence.

| **Location** | **MM** | **SM** |
| --- | --- | --- |
| Contig ID | k141_98629 | k141_111524 |
| Contig length (bp) | 3398 | 4353 |
| ARG ORFs (n) | 1 | 1 |
| CAT genus | Acinetobacter | Acinetobacter |
| CAT species | NA | NA |
| Terminal CAT rank | genus | genus |
| CAT genus support | 0.63 | 0.8 |
| Plasmid-like (geNomad) | TRUE | TRUE |
| geNomad plasmid score | 0.9678 | 0.818 |
| Virus-like (geNomad) | FALSE | FALSE |
| Provirus region | FALSE | FALSE |
| MGE ORFs (n) | 1 | 0 |
| MGE gene | 712_tnpA_EU850412.1 | NA |
| MGE class | transposase | NA |
| ARG-MGE same ORF | FALSE | FALSE |
| ARG-MGE overlapping ORFs | FALSE | FALSE |
| Nearest ARG-MGE distance (bp) | 2810 | NA |
| ARG-MGE within 5 kb | TRUE | FALSE |
| ResFinder genes | NA | NA |
| Context tier | Tier_2_mobile_context | Tier_2_mobile_context |

**Table S19D.** Genus-level HPB-list correspondence by site, relative to all canonical ACCs and to ACCs resolved to genus or species rank.

| **River** | **Site** | **All ACCs** | **ACCs resolved to genus or species** | **ACCs resolved to species** | **ACCs with genus-level HPB correspondence** | **% of all ACCs** | **% of genus/species-resolved ACCs** |
| --- | --- | --- | --- | --- | --- | --- | --- |
| Adyar | MM | 322 | 37 | 1 | 19 | 5.9 | 51.4 |
|  | SM | 787 | 221 | 71 | 107 | 13.6 | 48.4 |
| Musi | NH | 260 | 41 | 36 | 0 | 0 | 0 |
|  | NRH | 436 | 91 | 53 | 1 | 0.23 | 1.1 |
|  | MRH | 277 | 19 | 17 | 0 | 0 | 0 |
|  | SPH | 320 | 43 | 39 | 0 | 0 | 0 |
|  | All sites | 2402 | 452 | 217 | 127 | 5.29 | 28.1 |

**Table S19E.** ACCs resolved to species rank within HPB-listed genera, and their status on the HPB species list.

| **Site** | **SM** | **SM** |
| --- | --- | --- |
| CAT genus | *Pseudomonas* | *Pseudomonas* |
| CAT species | *Pseudomonas fluvialis* | *Pseudomonas mangrovi* |
| ACCs | 1 | 2 |
| On HPB species list | No | No |
